# Nab-paclitaxel fused with the *de novo* designed receptor binder exhibits enhanced tumor targeting and therapeutic efficacy

**DOI:** 10.64898/2026.01.28.702218

**Authors:** Yuanying Qian, Weikang Yan, Fan Xu, Yali Liu, Fabao Chen, Yue Lu, Zihan Zhang, Ao Gu, Ruobing Yu, Zhen Fang, Yang Yu, Maolan Li, Longxing Cao, Yingbin Liu, Yongning He

**Author notes:** Equal contributors (Yuanying Qian, Weikang Yan, Fan Xu, Yali Liu). Correspondence to: Y.H.

## Abstract

Chemotherapy has been widely used in cancer treatment, but most of the chemotherapeutic drugs rely mainly on passive accumulation due to lack of target specificity, which may lead to systemic toxicity and limited clinical utility. Recent advances in artificial intelligence-based protein design provide new opportunities for developing precision therapeutics. Here we modify the albumin-bound paclitaxel (Nab-PTX), one of the most widely used drugs in chemotherapies, by applying the fusion proteins of albumin with the *de novo* designed protein binders targeting human EGFR or HER2. The resulting particles, EGFRmb-Nab-PTX and HER2mb-Nab-PTX, retain the similar physicochemical properties of Nab-PTX while acquiring the receptor-specific binding capacities. The *in vitro* assays show that both binder-modified Nab-PTX particles have increased uptake and inhibitory effects significantly in the cancer cell lines with high receptor expression levels. Furthermore, the data from a xenograft model, including tumor growth, excised tumor analysis, organ histology, and fluorescence imaging, show that the binder-modified Nab-PTX enhances tumor accumulation, improves tumor suppression, and reduces off-target toxicity compared with the conventional Nab-PTX, suggesting that they may have promising clinical potential in cancer treatment. Overall, this strategy provides an adaptable modular platform for generating albumin-based chemotherapeutic drugs with target specificities, which can be readily customized with diverse target binders to enable precise cancer therapies.

## Introduction

Over a half century, chemotherapeutic agents remain indispensable in clinical treatment of cancer^1,2^. Despite their broad clinical use and antitumor efficacy, most chemotherapeutic agents do not have target specificities, which restricts effective drug delivery to cancer tissues and leads to systemic toxicity^2^. In the meantime, targeted therapies have emerged as an important complement to conventional chemotherapies in the past decades, enabling more precise elimination of cancer cells and minimizing damage to normal tissues. Among targeted strategies, small molecule inhibitors and antibodies are the two major calsses^3^. Small molecule inhibitors exhibit superior tissue penetration with better selectivity than conventional chemotherapeutic agents, yet their rapid clearance and off-target effects limit therapeutic benefit^4,5^. By contrast, antibodies offer high specificity but are hindered by poor tissue penetration^6^, potential immunogenicity, high costs and restricted target options^7^. To overcome these bottlenecks, antibody-drug conjugates (ADCs) are developed as promising strategies but also face challenges such as complex manufacturing, heterogeneous drug-to-antibody ratios, and dose-limiting toxicities^8–10^.

Recently, artificial intelligence (AI)-based technology has revolutionized protein structural prediction and design^11,12^ and creates new possibilities for next-generation therapeutics^13–15^. Over the past few years, *de novo* designed proteins, especially binders for various targets, have been applied in diverse contexts, including SARS-CoV-2 treatment^16,17^, cell signaling modulators^14,18^, cytokine mimetics^19^ , antibody-displaying nanocages for virus neutralization^20^, engineered protein nanoparticles for therapeutic delivery^21,22^, and targeted CAR-T therapy^23^. In these studies, the *de novo* designed protein binders usually have small molecular weights with high affinity and stability as well as low immunogenicity^13,17,19,23^, these features increase their potential in clinical use.

Among chemotherapeutic agents, paclitaxel is widely used against several types of cancer, including pancreatic, colorectal, gastric, breast and small cell lung cancer^24–27^. Paclitaxel can inhibit the depolymerization of microtubes and block cell cycles, which then lead to cell death^28,29^. However, the insolubility of paclitaxel in water limits its clinical application, which could be resolved by applying oil-based solvents such as Cremophor EL^30^, but the solvents may also increase the risk of adverse events, especially hypersensitivity reactions^31^. Albumin-bound paclitaxel (Nab-PTX) was then developed by encapsulating paclitaxel using human serum albumin (ALB) ^32^, which enhances permeability and retention effect (EPR)^28,33,34^ by interacting with albumin binding proteins such as glycoprotein 60 (gp60) and secreted protein acidic and rich in cysteine (SPARC) and also decreases the rate of anaphylaxis. Nevertheless, Nab-PTX still relies primarily on passive targeting, leading to paclitaxel accumulation in non-target tissues with serious adverse effects, such as peripheral neuropathy, neutropenia, anemia, fatigue and alopecia^24,35–37^. Therefore, lack of tumor specificity remains a major obstacle for Nab-PTX and limits its clinical utility.

Nab-PTX is included in the first-line medications for clinical treatment of late-stage pancreatic ductal adenocarcinoma (PDAC) and breast cancer. PDAC, currently among the most common causes of cancer-related death^38^, frequently overexpresses the epidermal growth factor receptor (EGFR), which has been closely implicated in tumor progression and regarded as an important therapeutic target^39–41^. Although several EGFR-targeted therapies have been developed for PDAC^42–45^, clinical efficacy is limiting due to the presence of various genetic heterogeneity and resistance mechanisms. Meanwhile, HER2 (also known as ERBB2), another member of the EGFR family, is also an important therapeutic target, especially for breast cancer^46–48^. Antibodies against HER2 such as trastuzumab, pertuzumab and the ADC T-DM1 have substantially improved overall survival in HER2-positive patients ^46,49^. However, similar to EGFR-targeted approaches, HER2-targeted therapies also face challenges such as off-target effects, insufficient efficacy in resistant tumors^50,51^. In contrast, chemotherapeutic drugs like paclitaxel provide potent cytotoxic activity but lack tumor specificity. Therefore, combining the strong antitumor effects of chemotherapies with receptor-specific targeting may provide a valuable strategy to enhance efficacy while minimizing adverse effects.

Here we modified Nab-PTX by fusing the computationally designed protein binders for EGFR and HER2 and generated modified Nab-PTX particles with target specificities for the receptors, thereby providing a modular platform that can be rationally adapted for receptor-directed paclitaxel delivery, and also offers a generic strategy to transform conventional chemotherapeutics into precise and targeted therapies.

## Results

### Design and expression of ALB fusion proteins targeting EGFR or HER2

A *de novo* designed protein binder EGFRmb that targets the N-terminal domain I of human EGFR with high affinity (Kd, 1.2 nM) has been reported previously (Fig. 1A)^14^. Following the similar binder design strategy^14^, we chose the N-terminal domain I of HER2 as the binding target for the HER2 binder (HER2mb) (Fig. 1B). After docking billions of amino acids at the HER2 binding surface, Rosetta molecular modeling suite was used to optimize the conformations of scaffolds and the critical binding residues. The designed candidates were filtered according to the parameters for binding affinity and solubility, and 8069 candidates were obtained. Then DNA oligos encoding the candidates were synthesized as a pool and displayed on the surface of yeast for high-throughput screening. After four rounds of screening using fluorescently labelled HER2 protein by flow cytometry (Fig. S1A), an HER2mb was identified (Fig. S1B), which contains three short alpha helices with a molecular weight of 10 kDa (Fig. 1B; Fig. S2B). HER2mb was then expressed and purified from *E. coli* (Fig. S2), and its interaction with HER2 was monitored by biolayer interferometry (BLI), which showed an affinity around 120 nM (Fig. 1G).

**Figure 1.**
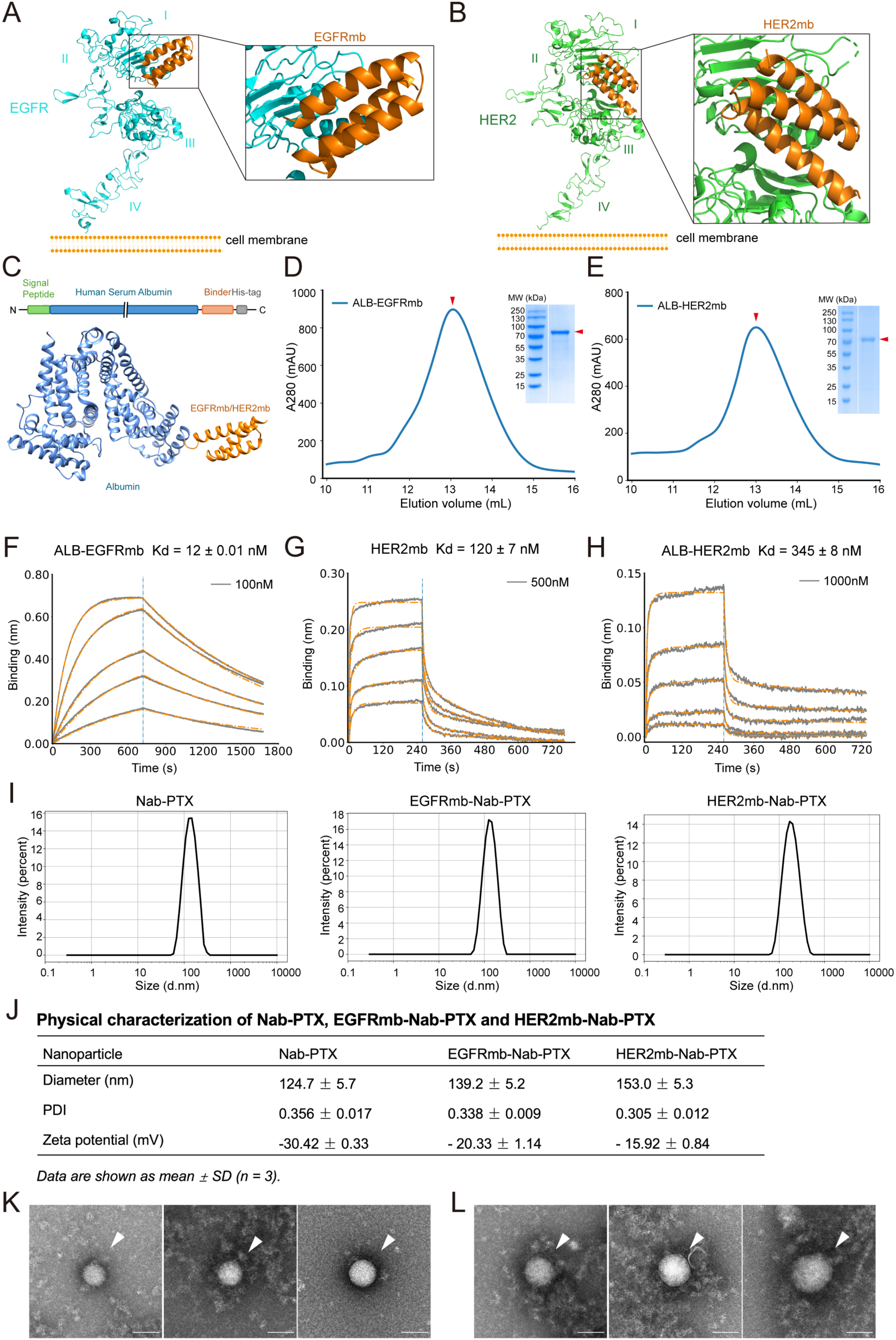
Design and characterization of the binder-modified Nab-PTX particles targeting EGFR and HER2. (A) A ribbon diagram model of EGFR ectodomain (cyan; PDB, 3NJP) bound with EGFRmb (orange) based on protein design and AlphaFold modeling. The domains (I-IV) of EGFR and the cell membrane are labeled. (B) A ribbon diagram model of HER2 ectodomain (green; PDB, 1N8Z) bound with HER2mb (orange) based on protein design and AlphaFold modeling. The domains (I-IV) of HER2 and the cell membrane are labeled. (C) The construct of ALB fused with EGFRmb or HER2mb (top). An AlphaFold model of the fusion protein is shown at the bottom. (D) The SEC profile and SDS-PAGE of purified fusion protein ALB-EGFRmb (red arrowheads). (E) The SEC profile and SDS-PAGE of purified fusion protein ALB-HER2mb (red arrowheads). (F) Binding of ALB-EGFRmb to EGFR ectodomain monitored by BLI. The highest concentration of ALB-EGFRmb is 100 nM, followed by serial two-fold dilutions. Experimental and fitted curves are shown in grey and orange, respectively. (G) Binding of HER2mb to HER2 ectodomain monitored by BLI. The highest concentration of HER2mb is 500 nM, followed by serial two-fold dilutions. Experimental and fitted curves are shown in grey and orange, respectively. (H) Binding of ALB-HER2mb to HER2 ectodomain by BLI. The highest concentration of ALB-HER2mb is 1000 nM, followed by serial two-fold dilutions. Experimental and fitted curves are shown in grey and orange, respectively. (I) Hydrodynamic diameter distributions of Nab-PTX, EGFRmb-Nab-PTX and HER2mb-Nab-PTX by DLS. (J) A summary of hydrodynamic diameter, polydispersity index (PDI), and zeta potential of the particles. (K) A gallery of EGFRmb-Nab-PTX particles (white arrowheads) by TEM (scale bar, 100 nm). (L) A gallery of HER2mb-Nab-PTX particles (white arrowheads) by TEM (scale bar, 100 nm).

The fusion proteins of ALB with the binders, named ALB-EGFRmb or ALB-HER2mb, were constructed by linking EGFRmb or HER2mb at the C-terminus of ALB (MW, 67 kDa) through a GSGS linker and a His-tag was included at the C-terminal end of the fusion proteins for purification (Fig. 1C). Since the binders are about one seventh of ALB in molecular weights, therefore may act as a small affinity tag for the receptors without affecting the properties of ALB, as shown in an AlphaFold model of the fusion protein (Fig. 1C). The fusion proteins were expressed and purified from insect cells with reasonable yields (Fig. 1D, E). The binding of the fusion proteins with EGFR or HER2 was evaluated by BLI (Fig. 1F, H). The data show that although the affinities of the binders for EGFR or HER2 were reduced after fusion with ALB, which is expected, the fusion proteins still retained nanomolar affinities with the receptors.

### Preparation and characterization of the modified Nab-PTX targeting EGFR or HER2

The nanoparticles of the modified albumin bound with paclitaxel targeting EGFR or HER2, named EGFRmb-Nab-PTX and HER2mb-Nab-PTX, were prepared by the emulsion-evaporation cross-linking method^52^ using ALB-EGFRmb or ALB-HER2mb instead of ALB, following a procedure similar to that for making conventional Nab-PTX. The particle size and zeta potential of the resulting particles were measured by dynamic light scattering (DLS). The data showed that the average hydrodynamic diameters of EGFRmb-Nab-PTX and HER2mb-Nab-PTX were around 139 nm and 153 nm, respectively, which were slightly bigger than that of Nab-PTX, which was around 123 nm (Fig. 1I, J). This is not surprising, as the fusion proteins are larger than ALB alone and may result in larger particles. Both EGFRmb-Nab-PTX and HER2mb-Nab-PTX exhibited comparable zeta potentials to non-modified Nab-PTX (Fig. 1J), indicating they had similar surface charge characteristics. Furthermore, the modified particles were also inspected under transmission electron microscopy (TEM), confirming the spherical shape and diameters around 100 nm and 120 nm for EGFRmb-Nab-PTX (Fig. 1K) and HER2mb-Nab-PTX (Fig. 1L), respectively.

The encapsulation efficiency (EE) of paclitaxel was also measured for EGFRmb-Nab-PTX, HER2mb-Nab-PTX and Nab-PTX, showing that EE values for all three particles were approximately 75% with small batch-to-batch variations, suggesting that the fusion proteins did not change the drug-loading capacity significantly. In addition, the binder-modified Nab-PTX particles were still active after storage at 4 °C for more than one month, suggesting these particles were quite stable.

### EGFRmb-Nab-PTX enhances the uptake of paclitaxel in cells with high EGFR expression levels

To assess the uptake of EGFRmb-Nab-PTX, a similar procedure was applied for preparing particles of EGFRmb-Nab-Cou-6 and Nab-Cou-6, where paclitaxel was replaced by a fluorescent probe coumarin-6 (Cou-6). Firstly, human kidney epithelial cells (HEK293T), which typically exhibit low basal EGFR expression^53^, were transfected with an mCherry-tagged EGFR expression plasmid. Flow cytometry data showed that the EGFR-overexpressed HEK293T cells internalized EGFRmb-Nab-Cou-6 much higher than Nab-Cou-6 (Fig. 2A), whereas the non-transfected cells showed almost no difference for EGFRmb-Nab-Cou-6 and Nab-Cou-6 (Fig. 2B), indicating enhanced targeting conferred by EGFRmb.

**Figure 2.**
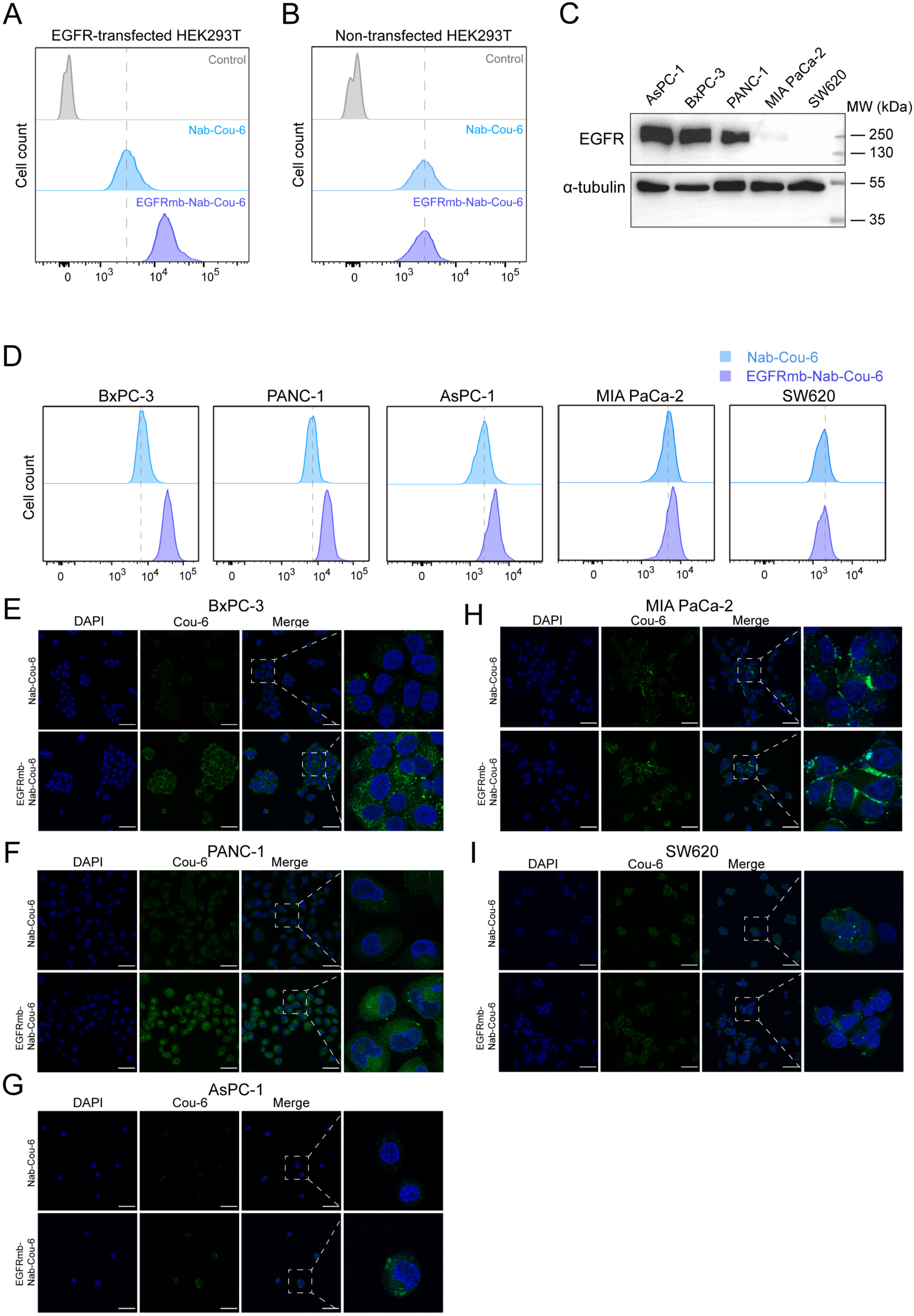
Cellular uptake of Nab-Cou-6 and EGFRmb-Nab-Cou-6. (A) Cellular uptake of Nab-Cou-6 or EGFRmb-Nab-Cou-6 in EGFR-transfected HEK293T cells by flow cytometry. (B) Cellular uptake of Nab-Cou-6 or EGFRmb-Nab-Cou-6 in non-transfected HEK293T cells by flow cytometry. (C) EGFR expression levels of the tumor cell lines (AsPC-1, BxPC-3, PANC-1, MIA PaCa-2, SW620) determined by western blot. Alpha-tubulin is applied as a loading control. (D) Cellular uptake of Nab-Cou-6 and EGFRmb-Nab-Cou-6 in the tumor cell lines by flow cytometry. (E-I) Fluorescence confocal images of the tumor cell lines treated with Nab-Cou-6 or EGFRmb-Nab-Cou-6. DAPI is applied for nucleus staining. Scale bar, 20 μm.

Then five cancer cell lines were applied, including four pancreatic cancer cell lines BxPC-3, PANC-1, AsPC-1 and MIA-PaCa2, and one colorectal cancer cell line SW620. The western blot assays showed that BxPC-3, PANC-1 and AsPC-1 had high EGFR expression, whereas MIA PaCa-2 and SW620 had low or almost undetectable expression of EGFR (Fig. 2C), which is consistent with the available protein expression data in the Human Protein Atlas^54^. The cell lines were incubated with EGFRmb-Nab-Cou-6 or Nab-Cou-6 separately and cellular uptake was monitored using flow cytometry (Fig. 2D), which showed that the cell lines with high EGFR expression, BxPC-3, PANC-1 and AsPC-1, internalized more EGFRmb-Nab-Cou-6 than Nab-Cou-6. By contrast, MIA PaCa-2, which has low EGFR expression, showed a slightly higher uptake of EGFRmb-Nab-Cou-6 than Nab-Cou-6, whereas SW620 exhibited similar uptake for both EGFRmb-Nab-Cou-6 and Nab-Cou-6. Then fluorescence confocal microscopy was applied to visualize intracellular uptake of the particles (Fig. 2E-I). Consistent with the flow cytometry data, EGFRmb-Nab-Cou-6 showed stronger intracellular fluorescence than Nab-Cou-6 in BxPC-3, PANC-1 and AsPC-1 (Fig. 2E-G), whereas no obvious difference was observed for both particles in MIA PaCa-2 and SW620 (Fig. 2H, I). These data suggest that uptake of EGFRmb-Nab-Cou-6 correlates with cellular EGFR expression, supporting the enhanced target specificity from the fusion protein.

### EGFRmb-Nab-PTX enhances antitumor efficacy in an EGFR expression-dependent manner

To investigate the antitumor efficacy of EGFRmb-Nab-PTX, the above cell lines were applied in cell viability assays. Since paclitaxel acts by disrupting microtubule dynamics during cell division, its potency may vary with the intrinsic proliferation rate of different cell lines, therefore growth rate (GR) inhibition metric GR_50_ was calculated as a normalized value to correct for the proliferation-dependent effects^55^, enabling a precise comparison of the differences in delivery between EGFRmb-Nab-PTX and Nab-PTX.

The cell lines were treated with either EGFRmb-Nab-PTX or Nab-PTX to assess drug responses in cell viability assays. For BxPC-3, PANC-1 and AsPC-1, which have high EGFR expression levels, GR_50_ values were significantly lower for EGFRmb-Nab-PTX than for Nab-PTX (Fig. 3A-D). By contrast, for the low EGFR expressing cells, MIA PaCa-2 and SW620, the GR_50_ differences between EGFRmb-Nab-PTX and Nab-PTX were narrowed down significantly (Fig. 3A, E, F). These results suggest that the enhanced efficacy of EGFRmb-Nab-PTX might be correlated with EGFR expression levels.

**Figure 3.**
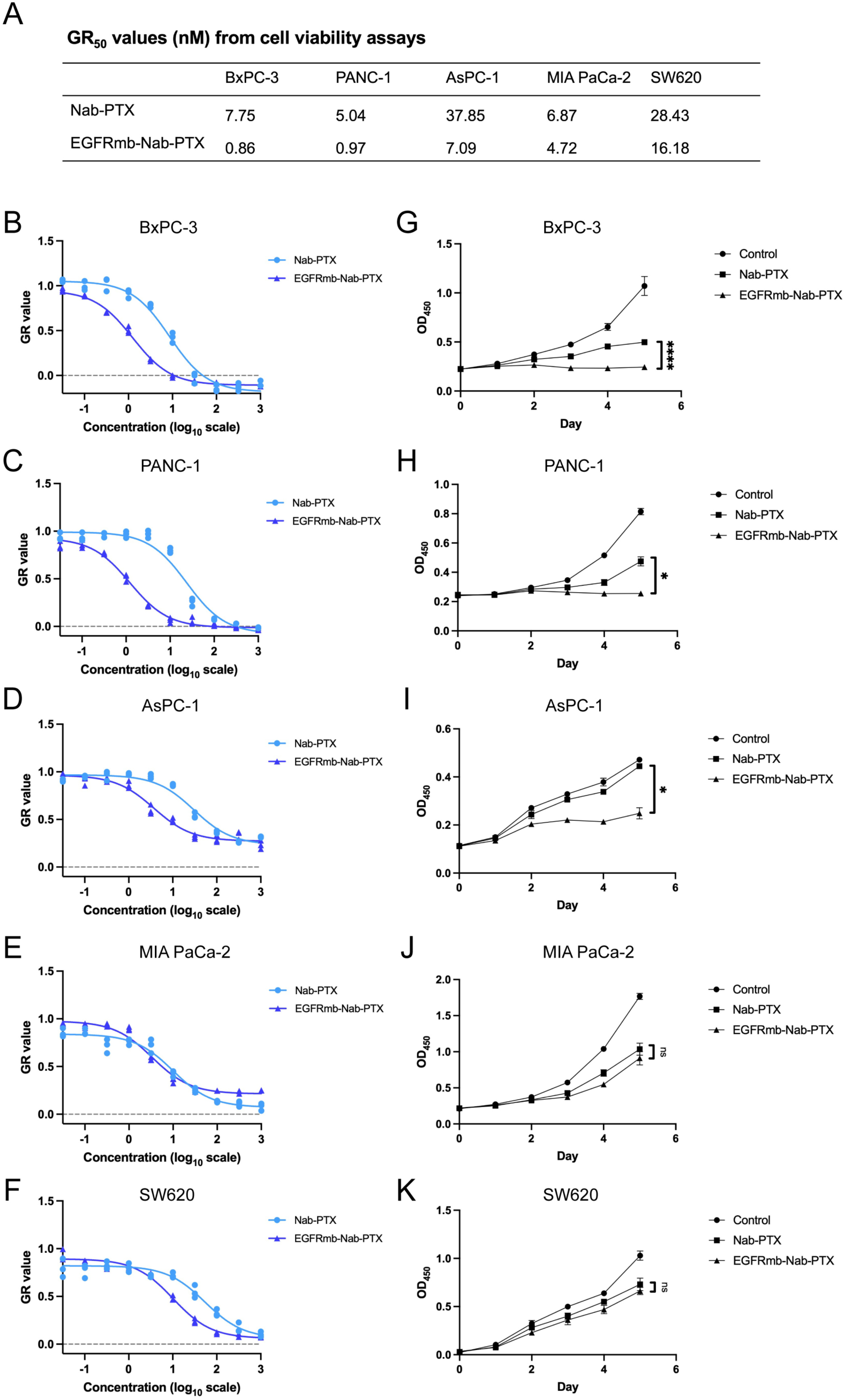
Cell viability and proliferation assays using Nab-PTX and EGFRmb-Nab-PTX. (A) GR_50_ values of the tumor cell lines after treatment with Nab-PTX or EGFRmb-Nab-PTX. (B-F) Cell viability assays for the tumor cell lines after treatment with Nab-PTX or EGFRmb-Nab-PTX based on GR metrics. Three replicates are performed for each concentration and all data points are plotted. (G-K) Cell proliferation assays for the tumor cell lines after treatment with Nab-PTX or EGFRmb-Nab-PTX. Each point represents the mean of three replicates and plotted as mean ± SD (n = 3). *p < 0.05; ****p < 0.0001; ns = not significant.

In parallel, cell proliferation assays were performed with EGFRmb-Nab-PTX or Nab-PTX at the same paclitaxel concentration. The results showed that EGFRmb-Nab-PTX could inhibit the proliferation of BxPC-3, PANC-1 and AsPC-1, which have high EGFR expression levels, significantly better than Nab-PTX (Fig. 3G-I), whereas the inhibitory effects of EGFRmb-Nab-PTX and Nab-PTX were similar for the low EGFR expression cells MIA PaCa-2 and SW620 (Fig. 3J, K). Notably, although AsPC-1 has a relatively high GR_50_ value in the selected cell lines (Fig. 3A), suggesting it is less sensitive to Nab-PTX, EGFRmb-Nab-PTX largely improved inhibitory effect for the cell line (Fig. 3I), supporting the enhanced efficacy is associated with EGFR expression levels.

Given that paclitaxel disrupts microtubule dynamics and induces cell cycle arrest at the G2/M phase, we analyzed the cell cycle distribution of the cell lines described above after treatment with EGFRmb-Nab-PTX or Nab-PTX (Fig. 4A-E). The results showed that the treatment of EGFRmb-Nab-PTX led to elongated G2/M phases in BxPC-3, PANC-1 and AsPC-1, which have high EGFR expression levels (Fig. 4A, B, C). By contrast, MIA PaCa-2 and SW620, which have low or negligible EGFR expression, exhibited similar G2/M phase distributions for EGFRmb-Nab-PTX and Nab-PTX (Fig. 4D, E). These results are consistent with the paclitaxel uptake, cell viability and proliferation data shown above.

**Figure 4.**
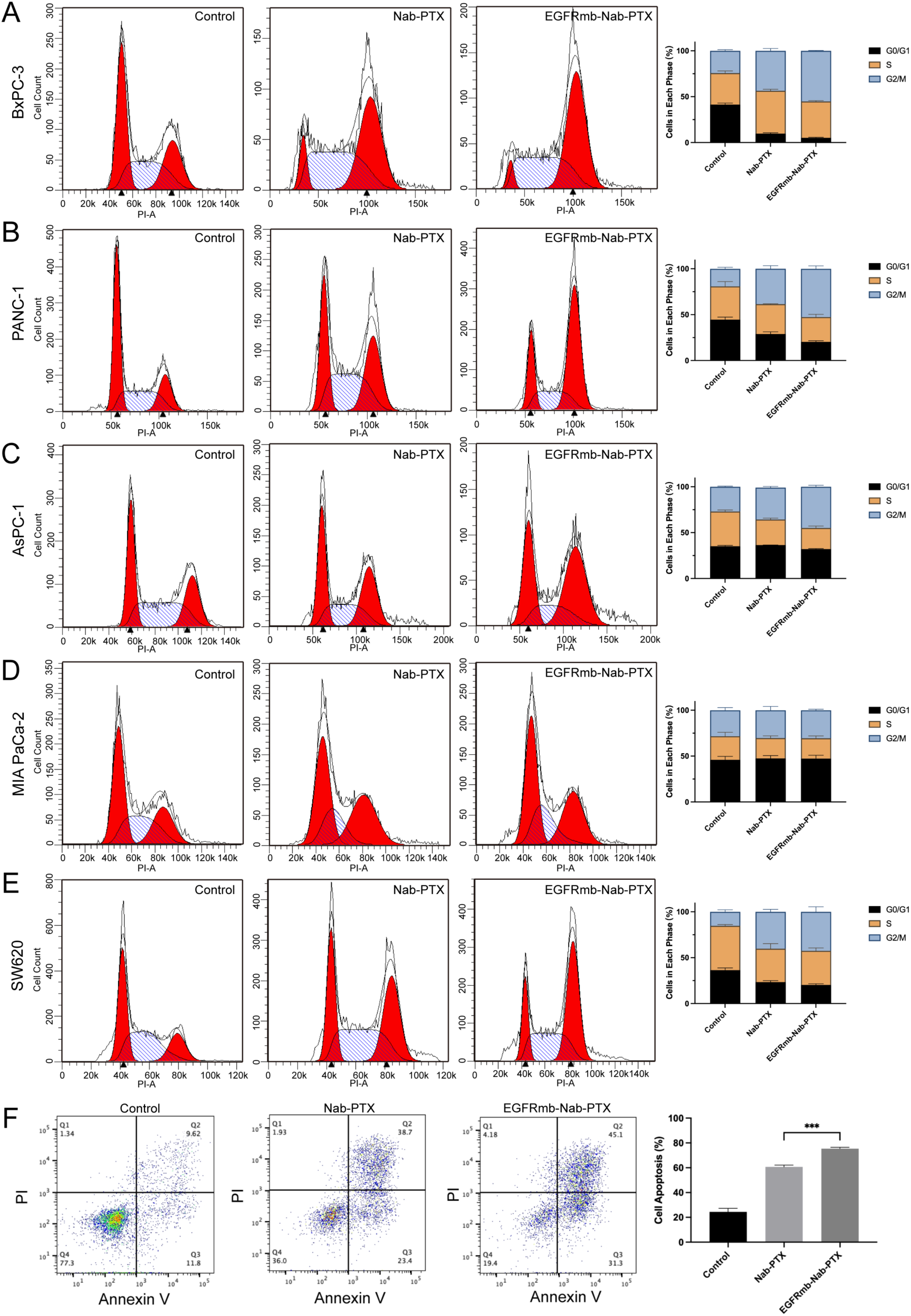
Cell cycle distribution and apoptosis analysis of the tumor cells treated with Nab-PTX and EGFRmb-Nab-PTX. (A-E) Cell cycle distribution in the tumor cells by flow cytometric analysis. Representative DNA content histograms of untreated cells and cells treated with Nab-PTX or EGFRmb-Nab-PTX are shown from left to right. The red areas represent the G0/G1 and G2/M phases, and the hatched blue areas represent the S phase. Quantitative summaries of the proportions of each phase are shown on the right. Data are presented as mean ± SD (n = 3). (F) Apoptotic ratio of BxPC-3 after treatment with Nab-PTX or EGFRmb-Nab-PTX by flow cytometric analysis. Untreated cells are applied as a control. Quantitative statistics are shown on the right. Data are presented as mean ± D (n = 3). ***p < 0.001.

It has been reported that Nab-PTX can trigger cell death through mitotic arrest and activate subsequent apoptotic pathways^56^. Here we monitored the apoptosis induced by EGFRmb-Nab-PTX or Nab-PTX using flow cytometry in BxPC-3, which has the highest EGFR expression among the cell lines examined above. Indeed, EGFRmb-Nab-PTX treatment resulted in a higher proportion of apoptotic cells (71.4%) than Nab-PTX (62.1%), supporting that EGFRmb-Nab-PTX enhanced intracellular accumulation of paclitaxel in cells with high EGFR expression.

### HER2mb-Nab-PTX enhances the uptake of paclitaxel in cells with high HER2 expression levels

Using the similar procedure for preparing EGFRmb-Nab-Cou-6, HER2mb-Nab-Cou-6 was made to evaluate the uptake of HER2mb-Nab-PTX in cell lines with different HER2 expression levels. Breast cancer cell lines SK-BR-3, BT-549 and MDA-MB-231 were selected, and western blot analysis showed that SK-BR-3 had high HER2 expression, while BT-549 and MDA-MB-231 had almost no HER2 expression (Fig. 5A), which is consistent with the available protein expression data in the Human Protein Atlas^54^. The cell lines were then incubated with HER2mb-Nab-Cou-6 or Nab-Cou-6 separately, and cellular uptake was monitored using both flow cytometry and fluorescence confocal microscopy.

**Figure 5.**
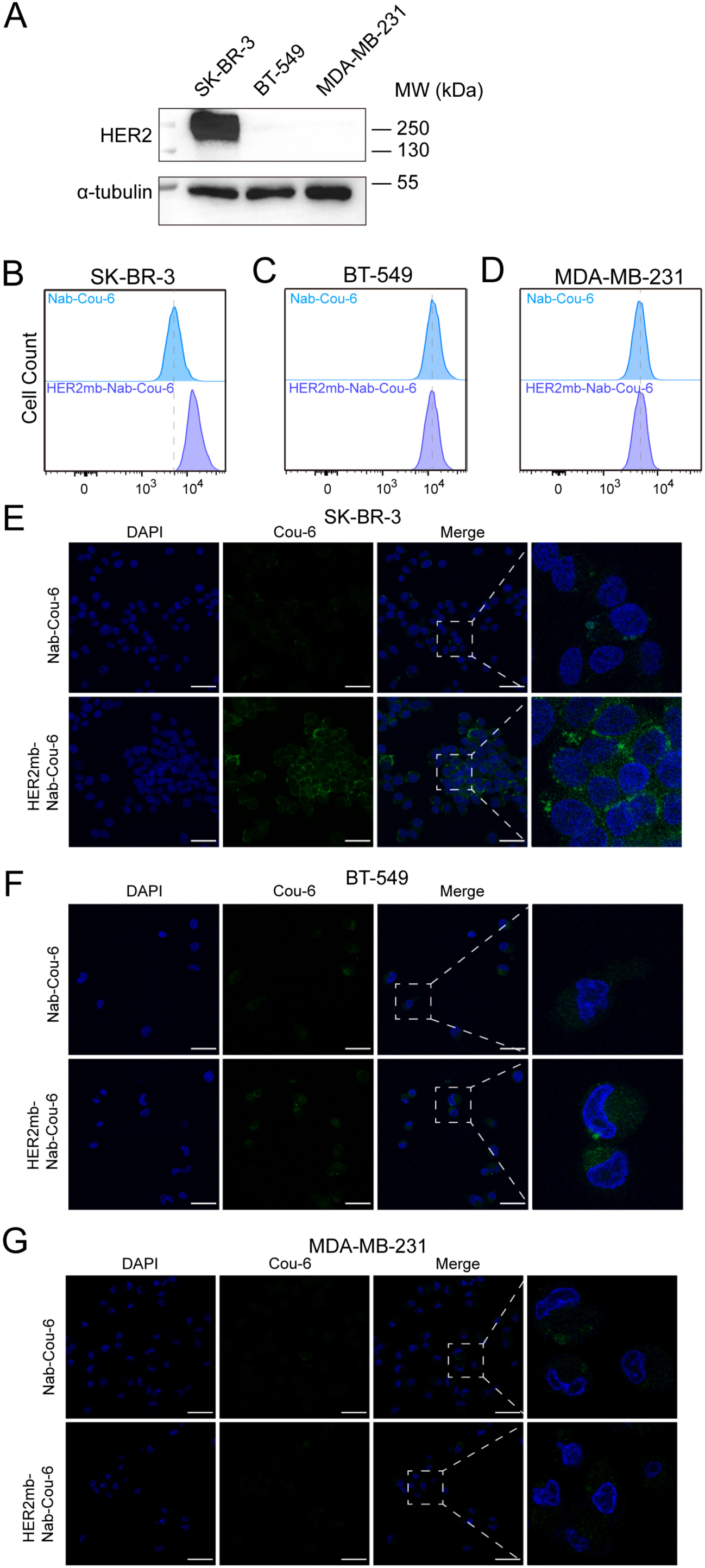
Cellular uptake of Nab-Cou-6 and HER2mb-Nab-Cou-6 in breast cancer cell lines. (A) HER2 expression levels in SKBR-3, BT-549 and MDA-MB-231 cells determined by western blot. Alpha-tubulin is applied as a loading control. (B-D) Cellular uptake of Nab-Cou-6 or HER2mb-Nab-Cou-6 in SKBR-3, BT-549 and MDA-MB-231 cells monitored by flow cytometry. (E-G) Fluorescence confocal images of the tumor cell lines treated with Nab-Cou-6 or HER2mb-Nab-Cou-6. DAPI is applied for nucleus staining. Scale bar, 20 μm.

Flow cytometry data showed that SK-BR-3 internalized HER2mb-Nab-Cou-6 more than Nab-Cou-6 (Fig. 5B), whereas BT-549 and MDA-MB-231 had similar uptake for HER2mb-Nab-Cou-6 and Nab-Cou-6 (Fig. 5C, D). The fluorescence confocal images also confirmed that SK-BR-3 cells exhibited stronger intracellular fluorescence after treatment with HER2mb-Nab-Cou-6 (Fig. 5E), and no obvious fluorescence was observed in BT-549 or MDA-MB-231 cells (Fig. 5F, G). These data suggest that HER2mb-Nab-Cou-6 was selectively internalized by cells due to HER2 expression.

### HER2mb-Nab-PTX enhances antitumor efficacy in a HER2 expression-dependent manner

To investigate the antitumor efficacy of HER2mb-Nab-PTX, the GR_50_ values were measured for SK-BR-3, BT-549 and MDA-MB-231after treatment with HER2mb-Nab-PTX or Nab-PTX (Fig. 6A-D). Analogous to the results from EGFRmb-Nab-PTX described above, HER2mb-Nab-PTX significantly reduced the GR_50_ for SK-BR-3 (Fig. 6A, B), but had similar values with Nab-PTX for BT-549 and MDA-MB-231 (Fig. 6A, C, D). In parallel, cell proliferation assays were performed to compare the treatments of HER2mb-Nab-PTX and Nab-PTX (Fig. 6E-G). The resulting data showed that HER2mb-Nab-PTX could significantly improve the inhibition of cell proliferation in SK-BR-3 (Fig. 6E), whereas the inhibitory effects were nearly indistinguishable with Nab-PTX in BT-549 and MDA-MB-231 (Fig. 6F, G).

**Figure 6.**
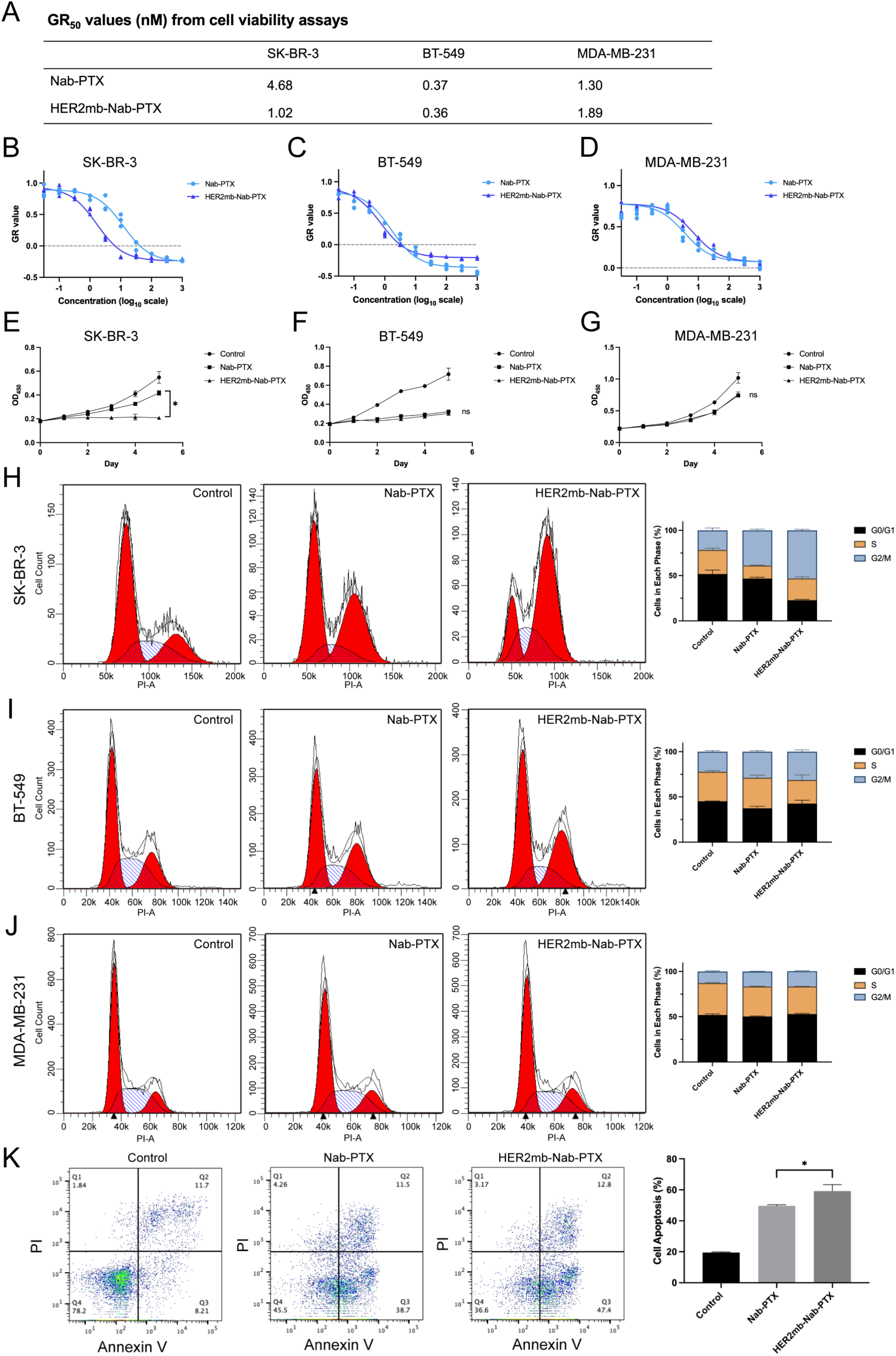
Cell viability, proliferation, cell cycle, and apoptosis analysis of the breast cancer cells treated with Nab-PTX and HER2mb-Nab-PTX. (A) GR_50_ values of the breast cell lines treated with Nab-PTX or HER2mb-Nab-PTX. (B-D) Cell viability assays for SK-BR-3, BT-549 and MDA-MB-231 after treatment with Nab-PTX or EGFRmb-Nab-PTX based on GR metrics. Three replicates are performed for each concentration and all data points are plotted. (E-G) Cell proliferation assays for SK-BR-3, BT-549 and MDA-MB-231 after treatment with Nab-PTX or EGFRmb-Nab-PTX. Each point represents the mean of three replicates and plotted as mean ± SD (n = 3). *p < 0.05, ns = not significant. (H-J) Cell cycle distribution in SK-BR-3, BT-549 and MDA-MB-231 cells by flow cytometric analysis. Representative DNA content histograms of untreated cells and cells treated with Nab-PTX or HER2mb-Nab-PTX are shown from left to right. The red areas represent the G0/G1 and G2/M phases, and the hatched blue areas represent the S phase. Quantitative summaries of the proportions of each phase are shown on the right. Data are presented as mean ± SD (n = 3). (K) Apoptotic ratio of SK-BR-3 cells after treatment with Nab-PTX or HER2mb-Nab-PTX by flow cytometric analysis. Untreated cells are applied as a control. Quantitative statistics are shown on the right. Data are presented as mean ± SD (n = 3). *p < 0.05.

Cell cycle distribution was also analyzed in the cell lines after treatment with HER2mb-Nab-PTX or Nab-PTX (Fig. 6H-J). For SK-BR-3, HER2mb-Nab-PTX treatment resulted in an elongated G2/M phase (53.20%) compared with Nab-PTX (38.88%) (Fig. 6H). By contrast, BT-549 and MDA-MB-231 cells showed comparable G2/M phase distributions after treatment with HER2mb-Nab-PTX or Nab-PTX (31.32% vs. 28.74% and 42.55% vs. 40.26%, respectively) (Fig. 6I, J). Apoptosis was also quantified in SK-BR-3 cells using flow cytometry (Fig. 6K), showing that Nab-PTX treatment induced apoptosis in 50.20% of cells, whereas HER2mb-Nab-PTX treatment increased this proportion to 60.20%, indicating enhanced intracellular accumulation of paclitaxel and stronger apoptotic induction in cells with high HER2 expression.

### EGFRmb-Nab-PTX improves antitumor efficacy and target specificity in vivo

To evaluate the antitumor efficacy and specificity *in vivo*, EGFRmb-Nab-PTX was applied in a subcutaneous xenograft model using BxPC-3 cells, and tumor growth was monitored following the treatment with EGFRmb-Nab-PTX or Nab-PTX (Fig. 7A). Compared with Nab-PTX, EGFRmb-Nab-PTX produced stronger inhibitory effects on tumor volumes (Fig. 7A) while exerting milder impacts on body weights (Fig. 7B). At the endpoint, tumors were excised, photographed and weighed. The tumors from the EGFRmb-Nab-PTX group were consistently smaller and lighter than those from the Nab-PTX group (Fig. 7C, D).

**Figure 7.**
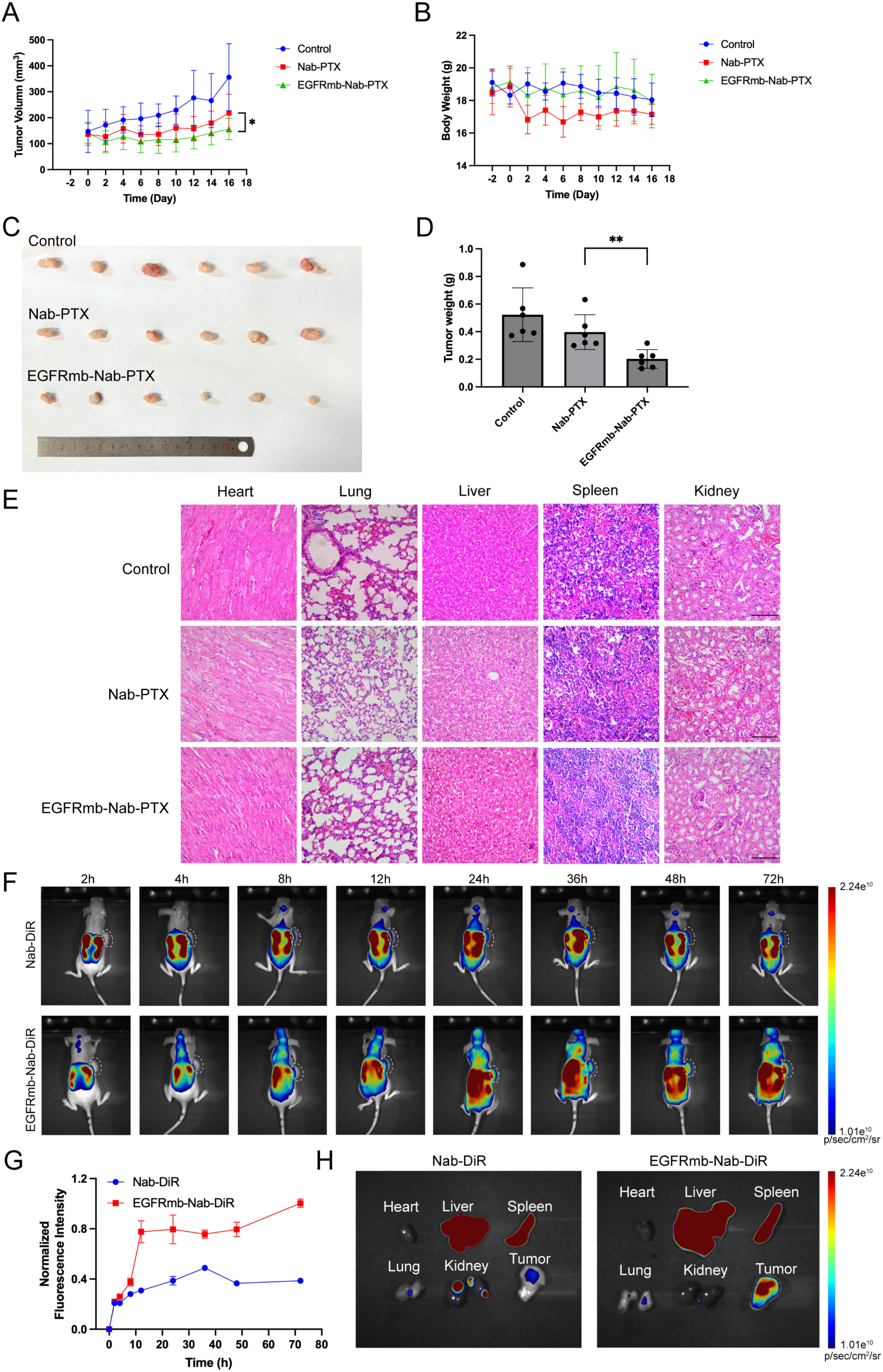
*In vivo* antitumor efficacy and biodistribution of Nab-PTX and EGFRmb-Nab-PTX. (A) Tumor growth curves of BxPC-3 tumor-bearing mice treated with saline (control), Nab-PTX, or EGFRmb-Nab-PTX (n=6). Data are presented as mean ± SD. *p < 0.05. (B) Body weight curves of the mice treated with saline (control), Nab-PTX, or EGFRmb-Nab-PTX (n=6). Data are presented as mean ± SD. (C) Representative images of the excised subcutaneous tumors from the control, the Nab-PTX, and the EGFRmb-Nab-PTX groups. (D) Average tumor weights from the three groups (n = 6). Data are presented as mean ± SD. **p < 0.01. (E) Representative H&E stained sections of the major organs (heart, liver, spleen, lung, and kidney) from the control, the Nab-PTX, and the EGFRmb-Nab-PTX groups. Scale bar, 100 μm. (F) *In vivo* fluorescence images of BxPC-3 tumor-bearing mice at the indicated time points after intraperitoneal injection of Nab-DiR or EGFRmb-Nab-DiR. Tumor locations are outlined by white dashed circles. (G) Fluorescence intensities of the tumors from the Nab-DiR and the EGFRmb-Nab-DiR group shown in (F). The highest fluorescence signal in tumor is normalized to one. Data are presented as mean ± SD (n=3). (H) *Ex vivo* fluorescence images of the major organs and tumors of the mice taken at 72 h from the Nab-DiR and EGFRmb-Nab-DiR group shown in (F).

To evaluate the potential systemic toxicity caused by the treatments, the major organs of the mice were collected for histological examination (Fig. 7E). Hematoxylin and eosin staining showed no evident pathological changes in heart, spleen or lung for both treated groups. Mild alterations were mainly observed in liver and kidney, which serve as primary organs for paclitaxel metabolism and clearance, and is in agreement with the previous reports showing that paclitaxel exposure is closely associated with hepatic functional impairment^57^. Notably, in the Nab-PTX group, hepatocellular vacuolar degeneration and limited inflammatory infiltration were observed in liver, accompanied by slight tubular epithelial swelling in kidney. By contrast, the EGFRmb-Nab-PTX group exhibited nearly normal hepatic and renal morphology, showing minimal histological alterations. Overall, these data suggest that EGFRmb-Nab-PTX achieves stronger tumor suppression with reduced off-target toxicity than Nab-PTX *in vivo*.

To compare the *in vivo* distribution of EGFRmb-Nab-PTX and Nab-PTX, DiR, a lipophilic fluorescent dye, was encapsulated in place of paclitaxel to prepare EGFRmb-Nab-DiR or Nab-DiR particles, which were administered by intraperitoneal injection. Serial *in vivo* fluorescence imaging was performed to visualize the distribution patterns of the particles (Fig. 7F). The tumor-associated fluorescence signal in the EGFRmb-Nab-DiR group gradually increased over time and was higher than that in the Nab-DiR group (Fig. 7F, G). At 72 h, the mice were sacrificed and the major organs together with the tumors were collected for *ex vivo* imaging (Fig. 7H). Among them, the tumor from the EGFRmb-Nab-DiR group exhibited markedly higher fluorescence intensity than that from the Nab-DiR group, indicating that EGFRmb modification enhances tumor specificity and promotes targeted accumulation in EGFR-positive tumors. Both groups showed strong fluorescence in liver and spleen, consistent with reticuloendothelial system uptake and hepatic clearance. In addition, the Nab-DiR group showed higher fluorescence signal in kidney than the EGFRmb-Nab-DiR group, which may correlate with the renal pathological alterations observed in the Nab-PTX treated mice (Fig. 7E, H). These results suggest that EGFRmb modification improves drug delivery selectivity and may also reduce off-target exposure.

## Discussion

Although targeted therapy and immunotherapy are developing rapidly over the past decades, chemotherapy still plays a fundamental role in cancer treatment. As a first-line drug for multiple cancers, including PDAC and breast cancers, Nab-PTX has shown substantial clinical efficacy, yet the lack of tumor specificity limits its therapeutic use. To increase tumor targeting, several albumin modification strategies have been published before, including passive-targeted modification with PEG or chitosan^58,59^, conjugation of antibodies or peptides to albumin nanoparticles^60,61^, physical adsorption or surface decoration with targeting ligands^62–65^ and fusion of albumin with functional domains such as collagen-binding domain^66^. While these methods have demonstrated efficacy enhancement, they often encounter limitations such as chemical heterogeneity, complex manufacturing processes, off-target accumulation or limited available protein domains.

Recently, protein design based on AI techniques has shown great potential in biomedical fields, as the *de novo* designed proteins usually have novel biological functions that cannot be achieved by natural proteins or small molecules. For example, protein design can generate specific binders for selected target proteins, thereby providing enormous opportunities in drug development and delivery. Here we utilize computationally *de novo* designed binders targeting EGFR or HER2 as representative examples for establishing a binder-albumin-based drug delivery platform. By fusing ALB with the designed binders, we generated modified Nab-PTX nanoparticles with target specificities for the receptors, which are supported by the subsequent *in vitro* and *in vivo* data. Therefore, this strategy may provide a broadly applicable platform for precise cancer therapies.

The characterization of the binder modified EGFRmb-Nab-PTX and HER2mb-Nab-PTX shows that both particles are similar to the unmodified Nab-PTX, with slightly increased particle sizes and similar paclitaxel encapsulation efficiencies.

Although binder affinities for the receptors are reduced after fusion with ALB, probably due to the steric hinderance from ALB, the binders utilized in this study still have nanomolar affinities for the receptors. At functional level, EGFRmb-Nab-PTX exhibits larger reductions in GR_50_ values than HER2mb-Nab-PTX. Moreover, the differences in paclitaxel uptake, growth inhibition and cell cycle arrest between EGFRmb-Nab-PTX and Nab-PTX are more evident than those between HER2mb-Nab-PTX and Nab-PTX, particularly in receptor high-expressing cells. These results suggest that binders with higher affinities might be more favorable for receptor targeting and drug delivery using this strategy. In addition, receptor abundance and accessibility, endocytic efficiency and nanoparticulophagy^67^ may also affect efficacy of the binder-modified Nab-PTX and require optimization for individual target receptors. Clinically, the therapeutic outcomes using this strategy need to be evaluated in the future, considering the tumor heterogeneity in target receptor expression in patients.

By expressing the fusion proteins of ALB with the binders, the strategy introduced in this study circumvents the challenges of chemical conjugation, such as random modification sites, structural perturbation or aggregation and variable conjugation efficiency, and can guarantee 100% modification of ALB. In addition, the fusion proteins expressed by insect cells usually have high yields and the whole procedure is cost effective. Compared with ADC where antibodies are used for targeting, the designed binders are not limited to the natural epitopes and can be tailored to novel binding sites or targets, thereby broadening the landscape of druggable molecules. However, although existing studies suggest low immunogenicity of the designed binders, further evaluation in clinical settings is necessary.

Taken together, our results show that fusing designed binders into albumin carriers provides a straightforward and generic method that can effectively transform conventional albumin-based chemotherapies into targeted treatments. Beyond EGFR and HER2, this modular platform can be extended to diverse tumor targets and offers a versatile strategy to enhance precision and efficacy of chemotherapies.

## Materials and Methods

### Cell lines and plasmids

HEK293T cells, pancreatic cancer cell lines AsPC-1, BxPC-3, PANC-1, MIA PaCa-2, the colorectal cancer cell line SW620 and the breast cancer cell line MDA-MB-231 were obtained from the Cell Bank of the Chinese Academy of Sciences (Shanghai, China). Breast cancer cell lines SK-BR-3, BT-549 were obtained from Procell Life Science & Technology Co., Ltd. (Wuhan, China). All cell lines were Short Tandem Repeat (STR) authenticated and tested mycoplasma free. All cell lines were all cultured at 37 °C under a humidified atmosphere containing 5% CO2. The pcDNA3 vector was purchased from General Biol (Anhui, China). A construct encoding human EGFR (NM_005228.5) with a C-terminal mCherry tag was sub-cloned into the pcDNA3.1 vectors for overexpressing EGFR.

### De novo design and screening of protein binder

The HER2 (PDB entry, 3H3B) targeting mini-protein binder was designed following a previously published workflow for *de novo* protein design^14^. Structural modeling and interface generation were performed using Rosetta FastRelax, RifDock, and RPXDock^68^, followed by sequence optimization with ProteinMPNN^69^ and structure validation with AlphaFold2. Top 8,069 designs based on the above metrics were selected and the oligonucleotide library encoding the designed binders were purchased from Agilent Technologies and amplified using KOD One^TM^ PCR Master Mix (TOYOBO, Japan). The amplified library was co-electroporated with the pETcon3 vector into the Saccharomyces cerevisiae EBY100 strain. Transformed yeast cells were cultured in SD-CAA medium. For protein expression, the cells were transferred into SG-CAA medium at a density of 0.3-0.5×10^7^ cells/mL and induced at 30°C and 225rpm for 12 hours. Following induction, the yeast cells were centrifuged at 3,000 ×g for 1min and washed with PBSF buffer (PBS supplemented with 0.1%(w/v) BSA). The cells were incubated with 20 μM biotinylated truncated HER2 protein for 30 min, washed with PBSF buffer, and subsequently labelled with anti-c-Myc FITC (Miltenyi Biotech, Germany) and streptavidin-PE (Thermo Fisher, USA) for 30 min. After washing and resuspending, the cells were analyzed and sorted by fluorescence-activated cell sorting (FACS). The final sorted pools were plated in SD-CAA plates, and individual clones were isolated and sequenced. The resulting sequence of HER2mb is: DAYGKASRLLHEAYKAYVRGDLETA AK LL KEGMKLLKKAGLENSQLYKSIKASLELVEKKIKEAK.

### Protein expression and purification

For expression and purification of HER2mb, the gene encoding HER2mb with an N-terminal 6× His tag was cloned into pET-28a(+) plasmid and transformed into chemically competent *E. coli* BL21(DE3) cells. Transformed BL21 were cultured in LB medium at 37°C and protein expression was induced with 0.5 mM isopropyl β-D-thiogalactoside (IPTG; Sangon, China) for 16 h at 37°C. Cells were harvested by centrifugation at 12,000 ×*g* for 10 min, resuspended in PBS buffer, and lysed by high-pressure homogenizer. After removal of cell debris, the supernatant was purified by Ni-NTA affinity chromatography (Smart Lifesciences, China), followed by size-exclusion chromatography (SEC) on a Superdex 75 Increase 10/300 GL column (Cytiva, Sweden) in PBS buffer (pH 7.4). The purity of the final protein was assessed by SDS-PAGE and Coomassie blue staining, the protein concentration was determined using a BCA Protein Quantification Kit (Yeasen, China). The yield of purified protein was around 1 mg per liter of culture.

For expression and purification of EGFR and HER2 ectodomain, the cDNA encoding the EGFR ectodomain (residue ID 25-525, UniProt ID P00533) or HER2 ectodomain (residue ID, 25-530, UniProt ID P04626) was cloned into the pTT5 vector, with an IL3 secretory signal peptide at the N-terminus and a 10×His-tag and a BirA biotinylation tag at the C-terminus. The plasmids were transfected into HEK293F cells using PEI 40000 (Yeasen, China) and cultured in Union 293 medium (Union Biotech, China) at 37°C with 8% CO₂ for 4 days. After culturing, the supernatant was centrifuged at 16,000 ×g for 60 minutes to remove cell debris. The proteins were purified using Ni Smart Beads (Smart Lifesciences, China), followed by SEC on a HiLoad 16/600 Superdex 200 pg column (Cytiva, Sweden) in 50mM Tris-HCl 150mM NaCl, pH 8.0. Protein concentrations of the purified protein were determined using a BCA assay, and biotinylation was performed with a BirA Biotin-Protein Ligase Kit (BirA500, Avidity, USA) according to the manuals. Excess biotin was further removed by SEC.

For expression and purification of albumin fusion proteins, the gene encoding human serum albumin (ALB) was cloned into the pFastBac1 vector (Invitrogen) with a C-terminal 6×His tag. The minibinder gene was inserted at the C-terminus of human albumin to generate ALB–binder fusion constructs. Recombinant plasmids were transformed into E. *coli* DH10Bac cells for bacmid generation following the Bac-to-Bac baculovirus system protocol (Invitrogen, USA). Positive colonies were selected, and bacmid DNA was extracted and verified by PCR. Verified bacmid was transfected into Sf9 cells to produce recombinant baculoviruses, which were then used to infect High Five (Hi5) cells for large-scale expression. Infected cells were cultured in ESF921 medium (Expression Systems, USA) at 27 °C for 3 days. The supernatants were collected, dialyzed against 25 mM Tris, 150 mM NaCl (pH 8.0), and purified by Ni–NTA affinity chromatography (Smart Lifesciences, China) followed by SEC on a HiLoad Superdex 200 10/300 pg column (Cytiva, Sweden). The purity of the final protein was assessed by SDS-PAGE and Coomassie blue staining, and the concentration was determined using BCA assays. The yields of purified fusion proteins were around 5-10 mg per liter of culture.

### Biolayer interferometry (BLI)

AVI-tagged EGFR and HER2 extracellular domains were first purified by SEC and exchanged into 50 mM Tris buffer (pH 8.0). Proteins were biotinylated using a BirA-RT Kit (Avidity, USA) at a concentration of 100 μM and incubated overnight at 4 °C, followed by desalting through SEC. The biotinylated EGFR and HER2 were then diluted to 25 μg/mL and 50 μg/mL, respectively, and immobilized onto streptavidin (SA) biosensors.

Purified ALB-EGFRmb, HER2mb, and ALB-HER2mb were used as analytes at highest concentrations of 100 nM, 500 nM, and 1000 nM, respectively, followed by serial two-fold dilutions. Binding buffer (PBS containing 0.5% BSA and 0.02% Tween-20) was applied for sensor equilibration, baseline stabilization, and sample dilution. Each assay was carried out with 200 μL of sample per well using an Octet RED96E system (ForteBio, USA). Data were analyzed with the Octet Data Analysis software provided by the manufacturer.

### Preparation of unmodified and modified Nab-PTX

Both unmodified and modified Nab-PTX were prepared using an emulsion– evaporation cross-linking method. Paclitaxel (Sagon, China) was dissolved in an organic solvent containing 95% acetone and 5% ethanol (v/v), while ALB (Aladdin Biochemical, China), ALB-EGFRmb, or ALB-HER2mb was dissolved in deionized water. The paclitaxel solution was added dropwise into the aqueous solution containing ALB or fusion proteins under continuous stirring with drug-to-protein ratio of 1:9 (w/w). Then the mixed solution was loaded onto a high-pressure homogenizer to obtain a stable emulsion. The organic solvents were the removed by rotary evaporation, and the resulting suspension was sterilized through a 0.22-µm filter and lyophilized for storage. The encapsulation efficiency of paclitaxel was determined by dissolving the lyophilized samples in methanol to precipitate denatured proteins, followed by spectrophotometric quantification of the supernatant at 228 nm^70,71^. Paclitaxel concentrations were calculated based on a pre-established standard curve. The encapsulation efficiency was calculated as the ratio of loaded paclitaxel to the total amount of paclitaxel used in the preparation.

Fluorescent nanoparticles, including Nab-Cou-6, EGFRmb-Nab-Cou-6, HER2mb-Nab-Cou-6, Nab-DiR, and EGFRmb-Nab-DiR, were prepared using the similar procedure where paclitaxel was substituted by coumarin-6 (MedChemExpress, China) or DiR (MedChemExpress, China).

### Dynamic light scattering

Hydrodynamic diameter, polydispersity index (PDI) and zeta potential of unmodified and modified Nab-PTX particles were measured using a Zetasizer Pro analyzer (Malvern Panalytical, UK). Samples were diluted 1:10 (v/v) in deionized water to obtain an appropriate scattering intensity for detection. Measurements were performed in triplicate at 25 °C, and results are presented as mean ± standard deviation (SD).

### Transmission electron microscopy

10 µL of EGFRmb-Nab-PTX or HER2mb-Nab-PTX was applied to a glow-discharged EM carbon grid and stained with 2% (w/v) uranyl acetate. The negatively stained EM grids were imaged on a Tecnai T12 microscope (FEI) operated at 120 kV and images were recorded at a nominal magnification of 67,000, using a 4k×4k Eagle CCD camera, corresponding to a pixel size of 1.74Å per pixel on the specimen.

### Western blot analysis

Total cellular proteins were extracted using RIPA lysis buffer (Yeasen, China) following the manuals. Equal amounts of protein were loaded onto SDS-PAGE (Bio-Rad Laboratories, USA) and transferred onto PVDF membranes. The membranes were blocked with 5% (w/v) nonfat milk in PBS containing 0.1% Tween 20 for 1h at room temperature. After washing three times with PBS, the membranes were incubated with HRP-conjugated anti-EGFR (Cell Signaling Technology, USA), anti-HER2 (Cell Signaling Technology, USA), or anti-α-tubulin antibodies (Abcam, UK). After washing, the signals were detected with the High-sig ECL western blotting substrate (Tanon, China).

### Flow cytometry

Cells were seeded in 6-well plates at a density of 3 × 10^5^ cells per well and incubated overnight at 37 °C, then washed with PBS and serum-starved for 2h. Subsequently, cells were incubated with or without different formulations of Nab-Cou-6 at a concentration of 25 μg/mL for 30 min at 37 °C. After the incubation, cells were washed and collected, subjected to centrifugation at 300 × *g* for 5 min.

Quantitative analysis of cellular uptake was performed using a BD Fortessa flow cytometer (BD Biosciences, USA), and 10,000 events were recorded for each sample. Untreated cells served as controls. Data were analyzed using FlowJo software (v 10.0).

### Confocal microscopy

Cells were seeded on glass coverslips placed in 12-well plates at a density of 5 × 10⁴ cells per well and incubated overnight at 37 °C. Cells were then treated with or without different formulations of Nab-Cou-6 following the same incubation procedures described above. After incubation, cells were washed three times with PBS, fixed with 4% paraformaldehyde, and permeabilized with 0.5% Triton X-100. Nuclei were stained using antifade mounting medium containing DAPI (Beyotime, China). Fluorescence confocal images were acquired using a Leica TCS SP8 confocal microscope. All experiments were performed in triplicate.

### Cell viability assays

Cells were seeded in 96-well plates at a density of 3 of 3 × 10³ cells per well and incubated overnight at 37 °C. Cells were then treated with or without different formulations of Nab-PTX at serial half-log (3.16-fold) dilutions. After 72 h of incubation, cell viability was measured using the Cell Counting Kit-8 (Yeasen, China) according to the manuals. Absorbance at 450 nm was recorded using a microplate reader (Thermo Fisher, USA). The absorbance values at the start and end of drug treatment were used for growth rate (GR) value calculation. GR_50_ values were finally calculated using the R package GR metrics^55^. Data were analyzed using GraphPad Prism.

### Cell growth assay

Cells were seeded in 96-well plates at a density of 1 × 10^3^ cells per well and incubated overnight at 37 °C. Cells were then treated with or without different formulations of Nab-PTX. Cell viability was measured daily from day 0 to day 5 using the Cell Counting Kit-8 (Yeasen, China) according to the manuals. Each condition was tested in triplicate. Data were analyzed using GraphPad Prism and presented as the mean ± SD (n=3). Differences between treatment groups were analyzed using two-way ANOVA, with *p* < 0.05 considered statistically significant.

### Cell cycle analysis

Cells were seeded in 6-well plates at a density of 3 × 10^5^ cells per well and incubated overnight at 37 °C. Upon reaching 80% confluence, the culture medium was replaced with serum-free medium. Cells were then treated with complete medium containing with or without different formulations of Nab-PTX at 37 °C for 12 h, with identical paclitaxel concentration for each cell line. Subsequently, cells were cultured in fresh complete medium for another 12 h, then washed with PBS and fixed with 70% cold ethanol overnight. Fixed cells were further collected, washed, and stained with 500 μL mix of RNase A and propidium iodide (PI) for 30 min at 37 °C in the dark. Stained samples were detected with FACS Fortessa flow cytometer (BD Biosciences), and the proportions of cell cycle phases were analyzed using ModFit LT Software. Statistical analysis was performed using GraphPad Prism and data were presented as the mean ± SD (n=3).

### Apoptosis analysis

Cells were seeded in 6-well plates at a density of 5 × 10^5^ per well and incubated for 24 h, followed by treatments with different formulations of Nab-PTX for 48 h.

The cells were then collected, washed twice and stained with annexin V and PI in the dark for 30 min at room temperature. Early (Annexin V^+^/PI^−^) and late (Annexin V^+^/PI^+^) apoptotic cells were identified by flow cytometry using FlowJo (v10.0), and their combined percentage was used to represent total apoptosis. Statistical analysis was performed using GraphPad Prism, and data were presented as the mean ± SD (n=3). Differences between treatment groups were analyzed using two-tailed unpaired *t*-tests, with *p* < 0.05 considered statistically significant.

### Animal model establishment

Pancreatic cancer cell line-derived xenograft (CDX) models were established for *in vivo* therapeutic evaluation. All animal procedures were approved by the Shanghai Medical Experimental Animal Care Commission and the Ethics Committee of Renji Hospital (Shanghai, China). BALB/c nude mice (5 weeks old, 15-20 g) were purchased from the Department of Experimental Animals, Renji Hospital (Shanghai, China). BxPC-3 cells were subcutaneously injected into the right axilla. All experiments were performed under specific pathogen-free (SPF) conditions. Mice were housed in groups of five per individually ventilated cage in a 12 h light/dark cycle, at 20 ± 2 °C and 55 ± 15% relative humidity, with ad libitum access to food and water.

### In vivo antitumor efficacy assay

When tumors reached 150–180 mm^3^, mice were randomly divided into three groups and treated with saline (control), Nab-PTX (5 mg/kg), or EGFRmb-nab-PTX (5 mg/kg). Treatments were performed once every three days for two weeks. Body weights and tumor volumes were measured every two days, and tumor volume was calculated as V = length × width^2^/2. At the end of the experiment, mice were sacrificed, then tumors were excised and weighed. Major organs (heart, liver, spleen, lung and kidney) were collected for histological examination. In brief, tissues were fixed in 4% paraformaldehyde, dehydrated, embedded in paraffin and sectioned. The sections were then stained with hematoxylin and eosin (H&E) following standard protocols. Images were acquired using a light microscope (Olympus BX43, Japan). Statistical analysis was performed using GraphPad Prism. Data are presented as the mean ± SD. Differences between treatment groups in tumor growth were analyzed using two-way ANOVA, and tumor weight differences were analyzed using two-tailed unpaired *t*-tests, with *p* < 0.05 considered statistically significant.

### In vivo biodistribution assay

BxPC-3 tumor–bearing nude mice received intraperitoneal administration with Nab-DiR or EGFRmb-Nab-DiR at a DiR dose of 0.5 mg/kg. Serial fluorescence imaging was performed at 2, 4, 8, 12, 24, 36, and 72 h post-administration using a Raycision Imaging 200 system. After that, the mice were sacrificed, and the tumors together with the major organs (heart, liver, spleen, lung, and kidney) were also collected for *ex vivo* fluorescence imaging using the system. The exposure time and other acquisition parameters were kept identical between the two groups. Tumor fluorescence was quantified using the onboard Raycision software with manually defined regions of interest (ROIs). Fluorescence signals were normalized to the maximum tumor intensity across all time points. Statistical analysis was performed using GraphPad Prism. Data are presented as the mean ±SD.

## Acknowledgements

We thank the Integrated Laser Microscopy and Electron Microscopy Systems at the National Facility for Protein Science in Shanghai (NFPS), Shanghai Advanced Research Institute, Chinese Academy of Sciences, China, for technical support. This work is supported by the State Key Laboratory of Systems Medicine for Cancer, Shanghai Cancer Institute, Renji Hospital, Shanghai Jiao Tong University School of Medicine. And we also thank the support from Innovative research team of high-level local universities in Shanghai (SHSMU-ZLCX20212601).

## Supplementary figures

**Figure S1.**
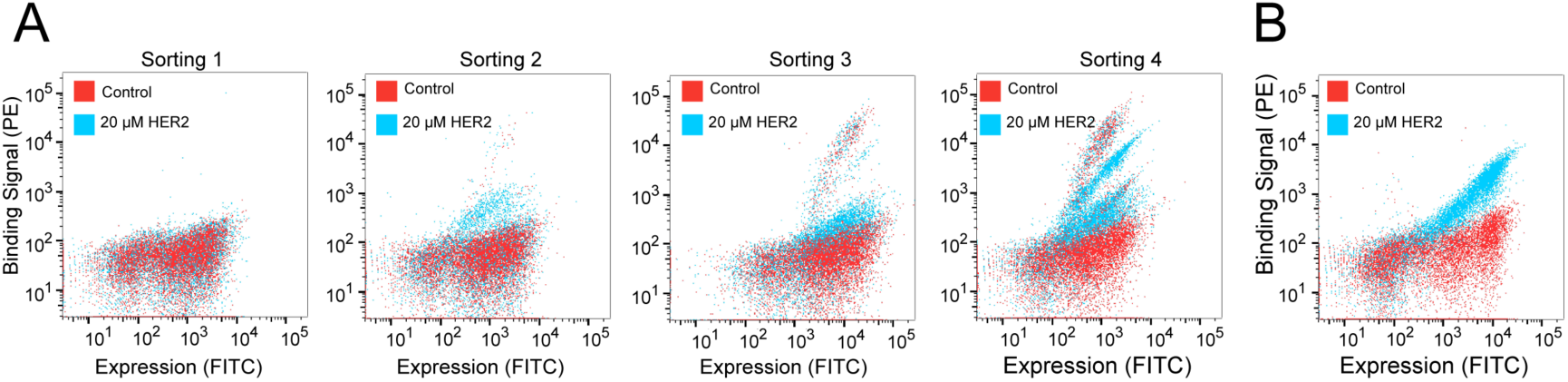
Screening and validation of the *de novo* designed binder targeting HER2 by flow cytometry. (A) Yeast surface display–based screening of HER2 binders over four rounds of cell sorting (Sorting 1–4). Yeast cells displaying the binder library were incubated with 20 μM recombinant HER2 protein. Binding signals were visualized as fluorescence intensity of target binding (y-axis) versus binder expression (x-axis). (B) Flow cytometric validation of the selected binder (HER2mb). Yeast cells displaying the single selected clone were incubated with 20 μM recombinant HER2 protein, and binding signals were detected as described above. Yeast cells displaying the library without recombinant HER2 protein incubation were applied as controls in (A) and (B).

**Figure S2.**
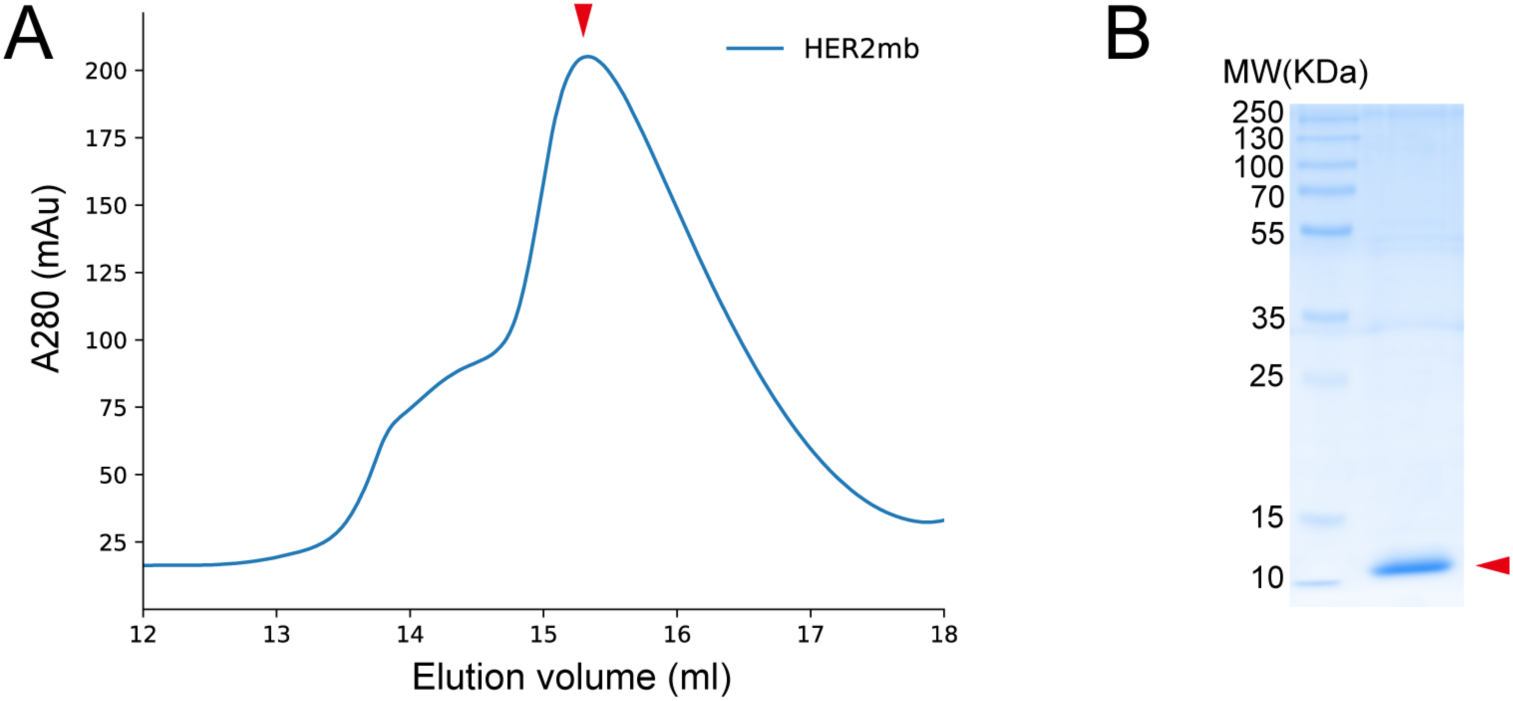
The SEC profile and SDS-PAGE of HER2mb. (A) The SEC profile of purified HER2mb. The elution peak corresponding to HER2mb is indicated by a red arrowhead. (B) The SDS-PAGE of purified HER2mb (red arrowhead).

## References

1. Cairns, J. (1985). The treatment of diseases and the war against cancer. Sci Am 253, 51–59. 10.1038/scientificamerican1185-51.

2. Anand, U., Dey, A., Chandel, A.K.S., Sanyal, R., Mishra, A., Pandey, D.K., De Falco, V., Upadhyay, A., Kandimalla, R., Chaudhary, A., et al. (2023). Cancer chemotherapy and beyond: Current status, drug candidates, associated risks and progress in targeted therapeutics. Genes Dis 10, 1367–1401. 10.1016/j.gendis.2022.02.007.

3. Bedard, P.L., Hyman, D.M., Davids, M.S., and Siu, L.L. (2020). Small molecules, big impact: 20 years of targeted therapy in oncology. Lancet 395, 1078–1088. 10.1016/S0140-6736(20)30164-1.

4. Lin, A., Giuliano, C.J., Palladino, A., John, K.M., Abramowicz, C., Yuan, M.L., Sausville, E.L., Lukow, D.A., Liu, L., Chait, A.R., et al. (2019). Off-target toxicity is a common mechanism of action of cancer drugs undergoing clinical trials. Sci Transl Med 11. 10.1126/scitranslmed.aaw8412.

5. Imai, K., and Takaoka, A. (2006). Comparing antibody and small-molecule therapies for cancer. Nat Rev Cancer 6, 714–727. 10.1038/nrc1913.

6. Bartelink, I.H., Jones, E.F., Shahidi-Latham, S.K., Lee, P.R.E., Zheng, Y., Vicini, P., van ’t Veer, L., Wolf, D., Iagaru, A., Kroetz, D.L., et al. (2019). Tumor Drug Penetration Measurements Could Be the Neglected Piece of the Personalized Cancer Treatment Puzzle. Clin Pharmacol Ther 106, 148–163. 10.1002/cpt.1211.

7. Sapra, P., and Shor, B. (2013). Monoclonal antibody-based therapies in cancer: advances and challenges. Pharmacol Ther 138, 452–469. 10.1016/j.pharmthera.2013.03.004.

8. Zhao, Z., Ukidve, A., Kim, J., and Mitragotri, S. (2020). Targeting Strategies for Tissue-Specific Drug Delivery. Cell 181, 151–167. 10.1016/j.cell.2020.02.001.

9. Colombo, R., and Rich, J.R. (2022). The therapeutic window of antibody drug conjugates: A dogma in need of revision. Cancer Cell 40, 1255–1263. 10.1016/j.ccell.2022.09.016.

10. Rubahamya, B., Dong, S., and Thurber, G.M. (2024). Clinical translation of antibody drug conjugate dosing in solid tumors from preclinical mouse data. Sci Adv 10, eadk1894. 10.1126/sciadv.adk1894.

11. Jumper, J., Evans, R., Pritzel, A., Green, T., Figurnov, M., Ronneberger, O., Tunyasuvunakool, K., Bates, R., Zidek, A., Potapenko, A., et al. (2021). Highly accurate protein structure prediction with AlphaFold. Nature 596, 583–589. 10.1038/s41586-021-03819-2.

12. Lin, Z., Akin, H., Rao, R., Hie, B., Zhu, Z., Lu, W., Smetanin, N., Verkuil, R., Kabeli, O., Shmueli, Y., et al. (2023). Evolutionary-scale prediction of atomic-level protein structure with a language model. Science 379, 1123–1130. 10.1126/science.ade2574.

13. Chevalier, A., Silva, D.A., Rocklin, G.J., Hicks, D.R., Vergara, R., Murapa, P., Bernard, S.M., Zhang, L., Lam, K.H., Yao, G., et al. (2017). Massively parallel de novo protein design for targeted therapeutics. Nature 550, 74–79. 10.1038/nature23912.

14. Cao, L., Coventry, B., Goreshnik, I., Huang, B., Sheffler, W., Park, J.S., Jude, K.M., Markovic, I., Kadam, R.U., Verschueren, K.H.G., et al. (2022). Design of protein-binding proteins from the target structure alone. Nature 605, 551–560. 10.1038/s41586-022-04654-9.

15. Huang, P.S., Boyken, S.E., and Baker, D. (2016). The coming of age of de novo protein design. Nature 537, 320–327. 10.1038/nature19946.

16. Cao, L., Goreshnik, I., Coventry, B., Case, J.B., Miller, L., Kozodoy, L., Chen, R.E., Carter, L., Walls, A.C., Park, Y.J., et al. (2020). De novo design of picomolar SARS-CoV-2 miniprotein inhibitors. Science 370, 426–431. 10.1126/science.abd9909.

17. Case, J.B., Chen, R.E., Cao, L., Ying, B., Winkler, E.S., Johnson, M., Goreshnik, I., Pham, M.N., Shrihari, S., Kafai, N.M., et al. (2021). Ultrapotent miniproteins targeting the SARS-CoV-2 receptor-binding domain protect against infection and disease. Cell Host Microbe 29, 1151–1161 e1155. 10.1016/j.chom.2021.06.008.

18. Park, J.S., Choi, J., Cao, L., Mohanty, J., Suzuki, Y., Park, A., Baker, D., Schlessinger, J., and Lee, S. (2022). Isoform-specific inhibition of FGFR signaling achieved by a de-novo-designed mini-protein. Cell Rep 41, 111545. 10.1016/j.celrep.2022.111545.

19. Silva, D.A., Yu, S., Ulge, U.Y., Spangler, J.B., Jude, K.M., Labao-Almeida, C., Ali, L.R., Quijano-Rubio, A., Ruterbusch, M., Leung, I., et al. (2019). De novo design of potent and selective mimics of IL-2 and IL-15. Nature 565, 186–191. 10.1038/s41586-018-0830-7.

20. Divine, R., Dang, H.V., Ueda, G., Fallas, J.A., Vulovic, I., Sheffler, W., Saini, S., Zhao, Y.T., Raj, I.X., Morawski, P.A., et al. (2021). Designed proteins assemble antibodies into modular nanocages. Science 372. 10.1126/science.abd9994.

21. Boyken, S.E., Benhaim, M.A., Busch, F., Jia, M., Bick, M.J., Choi, H., Klima, J.C., Chen, Z., Walkey, C., Mileant, A., et al. (2019). De novo design of tunable, pH-driven conformational changes. Science 364, 658–664. 10.1126/science.aav7897.

22. Eweje, F., Ibrahim, V., Shajii, A., Walsh, M.L., Ahmad, K., Alrefai, A., Miyasato, D., Davis, J.R., Ham, H., Li, K., et al. (2025). Self-assembling protein nanoparticles for cytosolic delivery of nucleic acids and proteins. Nat Biotechnol. 10.1038/s41587-025-02664-2.

23. Xia, Z., Jin, Q., Long, Z., He, Y., Liu, F., Sun, C., Liao, J., Wang, C., Wang, C., Zheng, J., et al. (2024). Targeting overexpressed antigens in glioblastoma via CAR T cells with computationally designed high-affinity protein binders. Nat Biomed Eng 8, 1634–1650. 10.1038/s41551-024-01258-8.

24. Von Hoff, D.D., Ervin, T., Arena, F.P., Chiorean, E.G., Infante, J., Moore, M., Seay, T., Tjulandin, S.A., Ma, W.W., Saleh, M.N., et al. (2013). Increased survival in pancreatic cancer with nab-paclitaxel plus gemcitabine. N Engl J Med 369, 1691–1703. 10.1056/NEJMoa1304369.

25. Brufsky, A. (2017). nab-Paclitaxel for the treatment of breast cancer: an update across treatment settings. Exp Hematol Oncol 6, 7. 10.1186/s40164-017-0066-5.

26. Hassan, M.S., Awasthi, N., Ponna, S., and von Holzen, U. (2023). Nab-Paclitaxel in the Treatment of Gastrointestinal Cancers-Improvements in Clinical Efficacy and Safety. Biomedicines 11. 10.3390/biomedicines11072000.

27. Zhu, L., and Chen, L. (2019). Progress in research on paclitaxel and tumor immunotherapy. Cell Mol Biol Lett 24, 40. 10.1186/s11658-019-0164-y.

28. Yardley, D.A. (2013). nab-Paclitaxel mechanisms of action and delivery. J Control Release 170, 365–372. 10.1016/j.jconrel.2013.05.041.

29. Long, B.H., and Fairchild, C.R. (1994). Paclitaxel inhibits progression of mitotic cells to G1 phase by interference with spindle formation without affecting other microtubule functions during anaphase and telephase. Cancer Res 54, 4355–4361.

30. ten Tije, A.J., Verweij, J., Loos, W.J., and Sparreboom, A. (2003). Pharmacological effects of formulation vehicles : implications for cancer chemotherapy. Clin Pharmacokinet 42, 665–685. 10.2165/00003088-200342070-00005.

31. Jones, S.E., Erban, J., Overmoyer, B., Budd, G.T., Hutchins, L., Lower, E., Laufman, L., Sundaram, S., Urba, W.J., Pritchard, K.I., et al. (2005). Randomized phase III study of docetaxel compared with paclitaxel in metastatic breast cancer. J Clin Oncol 23, 5542–5551. 10.1200/JCO.2005.02.027.

32. Ibrahim, N.K., Desai, N., Legha, S., Soon-Shiong, P., Theriault, R.L., Rivera, E., Esmaeli, B., Ring, S.E., Bedikian, A., Hortobagyi, G.N., and Ellerhorst, J.A. (2002). Phase I and pharmacokinetic study of ABI-007, a Cremophor-free, protein-stabilized, nanoparticle formulation of paclitaxel. Clin Cancer Res 8, 1038–1044.

33. Sleep, D. (2015). Albumin and its application in drug delivery. Expert Opin Drug Deliv 12, 793–812. 10.1517/17425247.2015.993313.

34. Chen, Q., Liang, C., Wang, C., and Liu, Z. (2015). An imagable and photothermal "Abraxane-like" nanodrug for combination cancer therapy to treat subcutaneous and metastatic breast tumors. Adv Mater 27, 903–910. 10.1002/adma.201404308.

35. Desai, N., Trieu, V., Yao, Z., Louie, L., Ci, S., Yang, A., Tao, C., De, T., Beals, B., Dykes, D., et al. (2006). Increased antitumor activity, intratumor paclitaxel concentrations, and endothelial cell transport of cremophor-free, albumin-bound paclitaxel, ABI-007, compared with cremophor-based paclitaxel. Clin Cancer Res 12, 1317–1324. 10.1158/1078-0432.CCR-05-1634.

36. Nyman, D.W., Campbell, K.J., Hersh, E., Long, K., Richardson, K., Trieu, V., Desai, N., Hawkins, M.J., and Von Hoff, D.D. (2005). Phase I and pharmacokinetics trial of ABI-007, a novel nanoparticle formulation of paclitaxel in patients with advanced nonhematologic malignancies. J Clin Oncol 23, 7785–7793. 10.1200/JCO.2004.00.6148.

37. Gradishar, W.J., Tjulandin, S., Davidson, N., Shaw, H., Desai, N., Bhar, P., Hawkins, M., and O’Shaughnessy, J. (2005). Phase III trial of nanoparticle albumin-bound paclitaxel compared with polyethylated castor oil-based paclitaxel in women with breast cancer. J Clin Oncol 23, 7794–7803. 10.1200/JCO.2005.04.937.

38. Siegel, R.L., Giaquinto, A.N., and Jemal, A. (2024). Cancer statistics, 2024. CA Cancer J Clin 74, 12–49. 10.3322/caac.21820.

39. Hynes, N.E., and Lane, H.A. (2005). ERBB receptors and cancer: the complexity of targeted inhibitors. Nat Rev Cancer 5, 341–354. 10.1038/nrc1609.

40. Barthel, S., Falcomata, C., Rad, R., Theis, F.J., and Saur, D. (2023). Single-cell profiling to explore pancreatic cancer heterogeneity, plasticity and response to therapy. Nat Cancer 4, 454–467. 10.1038/s43018-023-00526-x.

41. Moore, M.J., Goldstein, D., Hamm, J., Figer, A., Hecht, J.R., Gallinger, S., Au, H.J., Murawa, P., Walde, D., Wolff, R.A., et al. (2023). Erlotinib Plus Gemcitabine Compared With Gemcitabine Alone in Patients With Advanced Pancreatic Cancer: A Phase III Trial of the National Cancer Institute of Canada Clinical Trials Group. J Clin Oncol 41, 4714–4720. 10.1200/JCO.22.02770.

42. Blasco, M.T., Navas, C., Martin-Serrano, G., Grana-Castro, O., Lechuga, C.G., Martin-Diaz, L., Djurec, M., Li, J., Morales-Cacho, L., Esteban-Burgos, L., et al. (2019). Complete Regression of Advanced Pancreatic Ductal Adenocarcinomas upon Combined Inhibition of EGFR and C-RAF. Cancer Cell 35, 573–587 e576. 10.1016/j.ccell.2019.03.002.

43. Chiramel, J., Backen, A.C., Pihlak, R., Lamarca, A., Frizziero, M., Tariq, N.U., Hubner, R.A., Valle, J.W., Amir, E., and McNamara, M.G. (2017). Targeting the Epidermal Growth Factor Receptor in Addition to Chemotherapy in Patients with Advanced Pancreatic Cancer: A Systematic Review and Meta-Analysis. Int J Mol Sci 18. 10.3390/ijms18050909.

44. Momeny, M., Esmaeili, F., Hamzehlou, S., Yousefi, H., Javadikooshesh, S., Vahdatirad, V., Alishahi, Z., Mousavipak, S.H., Bashash, D., Dehpour, A.R., et al. (2019). The ERBB receptor inhibitor dacomitinib suppresses proliferation and invasion of pancreatic ductal adenocarcinoma cells. Cell Oncol (Dordr) 42, 491–504. 10.1007/s13402-019-00448-w.

45. Li, Z., Wang, M., Yao, X., Luo, W., Qu, Y., Yu, D., Li, X., Fang, J., and Huang, C. (2019). Development of a Novel EGFR-Targeting Antibody-Drug Conjugate for Pancreatic Cancer Therapy. Target Oncol 14, 93–105. 10.1007/s11523-018-0616-8.

46. Swain, S.M., Shastry, M., and Hamilton, E. (2023). Targeting HER2-positive breast cancer: advances and future directions. Nat Rev Drug Discov 22, 101–126. 10.1038/s41573-022-00579-0.

47. Meric-Bernstam, F., Johnson, A.M., Dumbrava, E.E.I., Raghav, K., Balaji, K., Bhatt, M., Murthy, R.K., Rodon, J., and Piha-Paul, S.A. (2019). Advances in HER2-Targeted Therapy: Novel Agents and Opportunities Beyond Breast and Gastric Cancer. Clin Cancer Res 25, 2033–2041. 10.1158/1078-0432.CCR-18-2275.

48. Oh, D.Y., and Bang, Y.J. (2020). HER2-targeted therapies - a role beyond breast cancer. Nat Rev Clin Oncol 17, 33–48. 10.1038/s41571-019-0268-3.

49. Fu, Z., Li, S., Han, S., Shi, C., and Zhang, Y. (2022). Antibody drug conjugate: the "biological missile" for targeted cancer therapy. Signal Transduct Target Ther 7, 93. 10.1038/s41392-022-00947-7.

50. Barok, M., Joensuu, H., and Isola, J. (2014). Trastuzumab emtansine: mechanisms of action and drug resistance. Breast Cancer Res 16, 209. 10.1186/bcr3621.

51. Pegram, M.D., Miles, D., Tsui, C.K., and Zong, Y. (2020). HER2-Overexpressing/Amplified Breast Cancer as a Testing Ground for Antibody-Drug Conjugate Drug Development in Solid Tumors. Clin Cancer Res 26, 775–786. 10.1158/1078-0432.CCR-18-1976.

52. Fu, Q., Sun, J., Zhang, W., Sui, X., Yan, Z., and He, Z. (2009). Nanoparticle albumin-bound (NAB) technology is a promising method for anti-cancer drug delivery. Recent Pat Anticancer Drug Discov 4, 262–272. 10.2174/157489209789206869.

53. Kancha, R.K., von Bubnoff, N., Peschel, C., and Duyster, J. (2009). Functional analysis of epidermal growth factor receptor (EGFR) mutations and potential implications for EGFR targeted therapy. Clin Cancer Res 15, 460–467. 10.1158/1078-0432.CCR-08-1757.

54. Uhlen, M., Fagerberg, L., Hallstrom, B.M., Lindskog, C., Oksvold, P., Mardinoglu, A., Sivertsson, A., Kampf, C., Sjostedt, E., Asplund, A., et al. (2015). Proteomics. Tissue-based map of the human proteome. Science 347, 1260419. 10.1126/science.1260419.

55. Hafner, M., Niepel, M., Chung, M., and Sorger, P.K. (2016). Growth rate inhibition metrics correct for confounders in measuring sensitivity to cancer drugs. Nat Methods 13, 521–527. 10.1038/nmeth.3853.

56. Khing, T.M., Choi, W.S., Kim, D.M., Po, W.W., Thein, W., Shin, C.Y., and Sohn, U.D. (2021). The effect of paclitaxel on apoptosis, autophagy and mitotic catastrophe in AGS cells. Sci Rep 11, 23490. 10.1038/s41598-021-02503-9.

57. Schmidt, M., Vernooij, R., van Nuland, M., Smeijsters, E., Devriese, L., Mohammad, N.H., Hermens, T., Stammers, J., Swart, C., Egberts, T., et al. (2024). Impaired liver function: effect on paclitaxel toxicity, dose modifications and overall survival. BMC Cancer 24, 1553. 10.1186/s12885-024-13330-2.

58. Fahrlander, E., Schelhaas, S., Jacobs, A.H., and Langer, K. (2015). PEGylated human serum albumin (HSA) nanoparticles: preparation, characterization and quantification of the PEGylation extent. Nanotechnology 26, 145103. 10.1088/0957-4484/26/14/145103.

59. Esfandyari-Manesh, M., Mohammadi, A., Atyabi, F., Nabavi, S.M., Ebrahimi, S.M., Shahmoradi, E., Varnamkhasti, B.S., Ghahremani, M.H., and Dinarvand, R. (2016). Specific targeting delivery to MUC1 overexpressing tumors by albumin-chitosan nanoparticles conjugated to DNA aptamer. Int J Pharm 515, 607–615. 10.1016/j.ijpharm.2016.10.066.

60. Kouchakzadeh, H., Shojaosadati, S.A., Tahmasebi, F., and Shokri, F. (2013). Optimization of an anti-HER2 monoclonal antibody targeted delivery system using PEGylated human serum albumin nanoparticles. Int J Pharm 447, 62–69. 10.1016/j.ijpharm.2013.02.043.

61. Wagner, S., Rothweiler, F., Anhorn, M.G., Sauer, D., Riemann, I., Weiss, E.C., Katsen-Globa, A., Michaelis, M., Cinatl, J., Jr., Schwartz, D., et al. (2010). Enhanced drug targeting by attachment of an anti alphav integrin antibody to doxorubicin loaded human serum albumin nanoparticles. Biomaterials 31, 2388–2398. 10.1016/j.biomaterials.2009.11.093.

62. Brindisi, M., Curcio, M., Frattaruolo, L., Cirillo, G., Leggio, A., Rago, V., Nicoletta, F.P., Cappello, A.R., and Iemma, F. (2022). CD44-targeted nanoparticles with GSH-responsive activity as powerful therapeutic agents against breast cancer. Int J Biol Macromol 221, 1491–1503. 10.1016/j.ijbiomac.2022.09.157.

63. Thao le, Q., Byeon, H.J., Lee, C., Lee, S., Lee, E.S., Choi, Y.W., Choi, H.G., Park, E.S., Lee, K.C., and Youn, Y.S. (2016). Doxorubicin-Bound Albumin Nanoparticles Containing a TRAIL Protein for Targeted Treatment of Colon Cancer. Pharm Res 33, 615–626. 10.1007/s11095-015-1814-z.

64. Min, S.Y., Byeon, H.J., Lee, C., Seo, J., Lee, E.S., Shin, B.S., Choi, H.G., Lee, K.C., and Youn, Y.S. (2015). Facile one-pot formulation of TRAIL-embedded paclitaxel-bound albumin nanoparticles for the treatment of pancreatic cancer. Int J Pharm 494, 506–515. 10.1016/j.ijpharm.2015.08.055.

65. Wang, X., Tian, L., Li, Y., Yao, W., Zhu, J., Zhou, H., Chen, G., Chen, T., Liu, Z., Tan, W., and Yang, Y. (2025). Universal Albumin Drugs-Cored Spherical Nucleic Acid (ad-SNA) Platform for Targeted Drug Delivery. Angew Chem Int Ed Engl 64, e202421949. 10.1002/anie.202421949.

66. Sasaki, K., Ishihara, J., Ishihara, A., Miura, R., Mansurov, A., Fukunaga, K., and Hubbell, J.A. (2019). Engineered collagen-binding serum albumin as a drug conjugate carrier for cancer therapy. Sci Adv 5, eaaw6081. 10.1126/sciadv.aaw6081.

67. Lin, Y.W., Lin, T.T., Chen, C.H., Wang, R.H., Lin, Y.H., Tseng, T.Y., Zhuang, Y.J., Tang, S.Y., Lin, Y.C., Pang, J.Y., et al. (2023). Enhancing Efficacy of Albumin-Bound Paclitaxel for Human Lung and Colorectal Cancers through Autophagy Receptor Sequestosome 1 (SQSTM1)/p62- Mediated Nanodrug Delivery and Cancer therapy. ACS Nano 17, 19033–19051. 10.1021/acsnano.3c04739.

68. Sheffler, W., Yang, E.C., Dowling, Q., Hsia, Y., Fries, C.N., Stanislaw, J., Langowski, M.D., Brandys, M., Li, Z., Skotheim, R., et al. (2023). Fast and versatile sequence-independent protein docking for nanomaterials design using RPXDock. PLoS Comput Biol 19, e1010680. 10.1371/journal.pcbi.1010680.

69. Dauparas, J., Anishchenko, I., Bennett, N., Bai, H., Ragotte, R.J., Milles, L.F., Wicky, B.I.M., Courbet, A., de Haas, R.J., Bethel, N., et al. (2022). Robust deep learning-based protein sequence design using ProteinMPNN. Science 378, 49–56. 10.1126/science.add2187.

70. Esfandyari-Manesh, M., Mostafavi, S.H., Majidi, R.F., Koopaei, M.N., Ravari, N.S., Amini, M., Darvishi, B., Ostad, S.N., Atyabi, F., and Dinarvand, R. (2015). Improved anticancer delivery of paclitaxel by albumin surface modification of PLGA nanoparticles. Daru 23, 28. 10.1186/s40199-015-0107-8.

71. Deng, W., Qiu, J., Wang, S., Yuan, Z., Jia, Y., Tan, H., Lu, J., and Zheng, R. (2018). Development of biocompatible and VEGF-targeted paclitaxel nanodrugs on albumin and graphene oxide dual-carrier for photothermal-triggered drug delivery in vitro and in vivo. Int J Nanomedicine 13, 439–453. 10.2147/IJN.S150977.

